# Bayesian adaptive experimental design for efficient microbial genome-wide association studies

**DOI:** 10.64898/2026.08.26.747358

**Authors:** David Helekal, Sofia O. P. Blomqvist, Aditi Mukherjee, Bailey Bowcutt, Samantha G. Palace, Yonatan H. Grad

## Abstract

Bacterial genome-wide association studies (GWAS) offer a powerful approach to identify the genetic basis of a trait measured in a set of sequenced isolates. As the number of sequenced isolates has grown, the limiting factor for GWAS has become phenotyping enough isolates to achieve statistical power. To overcome the need for large-scale phenotyping, we developed Bayesian Adaptive Sequential Sampling GWAS (BASS-GWAS), which couples Bayesian adaptive experimental design with a sparse regression model to select maximally informative isolates for phenotypic testing. BASS-GWAS efficiently recovered causal loci for three antimicrobial resistance traits in *Neisseria gonorrhoeae*, requiring many fewer phenotyped isolates than random sampling. We applied BASS-GWAS to discover variants enabling *gyrB*^D429N^-dependent cross-resistance to the novel topoisomerase inhibitors zoliflodacin and gepotidacin. After phenotyping fewer than 30 isolates, we identified and then validated both *parC*^D86N^ and a *gyrA-parE*-based pathway as enabling cross-resistance. BASS-GWAS provides a practical and statistically principled solution for efficient bacterial GWAS.

## Introduction

Bacterial genome-wide association studies (GWAS) offer a powerful strategy for identifying genetic variants that explain phenotypes of interest. Until recently, the limiting resource for bacterial GWAS was genome sequencing. Studies mostly relied on collections of isolates phenotyped as part of clinical evaluation or surveillance programs, and many isolates were sequenced in the hope that the collection would encompass sufficient genetic diversity to detect an association. This approach proved fruitful for phenotypes such as antimicrobial susceptibility and invasiveness, enabling a series of studies that identified and experimentally validated genetic determinants of these phenotypes [1–7].

These efforts resulted in large collections of sequenced bacterial isolates and created the possibility of using GWAS to identify the genetic basis of any measurable trait. As phenotyping is now the limiting step, a key question is which and how many isolates to assay. GWAS of randomly selected isolates may require hundreds to thousands of isolates for sufficient statistical power [2, 6, 8–12], even for traits with large effect sizes [13]. Experiments at this scale are prohibitive for many phenotypes. A strategy is needed to optimize experimental efficiency in phenotyping for GWAS, given a starting library of sequenced bacterial genomes.

To address this need, we developed Bayesian Adaptive Sequential Sampling GWAS (BASS-GWAS) that incorporates Bayesian Adaptive Design (BAD) and a whole-genome regression model. BAD is a framework for selecting experiments that maximize cumulative information gained [14–16], enabling highly efficient returns from resource-intensive experiments [15–21].

We used BASS-GWAS to identify the genetic basis of cross-resistance between zoliflodacin and gepotidacin, each a first-in-class antimicrobial recently approved for the treatment of *Neisseria gonorrhoeae* [22]. Although both drugs target topoisomerase complexes, initial characterization of resistant isolates pointed to distinct resistance mutations, primarily *gyrB*^D429N^ for zoliflodacin [23] and *gyrA*^A92T^ in combination with pre-existing *parC*^D86N^ for gepotidacin [24]. Unexpectedly, recent work demonstrated the existence of strain-specific cross-resistance between these drugs: the introduction of *gyrB*^D429N^ caused a 32-fold increase in gepotidacin resistance in two isolates from distinct genetic backgrounds, but did not alter gepotidacin resistance in several other isolates [25]. Understanding the genetic basis of this cross-resistance is important to inform treatment and surveillance strategies as zoliflodacin and gepotidacin enter clinical use. However, the set of isolates tested in the recent study was too small for GWAS, and targeted mutagenesis is too labor-intensive to test hundreds of randomly selected isolates. We therefore applied BASS-GWAS to identify the genetic determinants of *gyrB*^D429N^–mediated zoliflodacin and gepotidacin cross-resistance in *N. gonorrhoeae*.

## Results

### Overview of the Approach

BASS-GWAS employs iterative phenotyping of small groups of isolates (Figure 1). Starting with a sequenced collection of isolates available for experimental manipulation (Figure 1a), BASS-GWAS uses phenotypic information (Figure 1b) to update the posterior distribution of the GWAS model (Figure 1c) that estimates the effects of all genetic variants, evaluates how phenotypic information for unobserved strains would reduce uncertainty in these estimates, and employs BAD to identify a small batch of strains (4-20) that maximizes the expected information gain (EIG; [14–16]) for the next batch of assays (Figure 1d). This process is then iterated for a fixed number of iterations or until one or more candidate variants are identified with sufficient confidence to justify validation experiments. This approach improves efficiency of mapping genotype/phenotype associations by focusing on lineages with observed phenotypic variation while also testing unexplored regions of the phylogeny. We implemented BASS-GWAS with real-world experimental workflows in mind, such that (a) a batch of strains can be optimized conditional on including strains of interest, and (b) assays that fail can be censored from a batch, enabling removal of strains that prove nonviable or otherwise impossible to assay.

**Figure 1:**
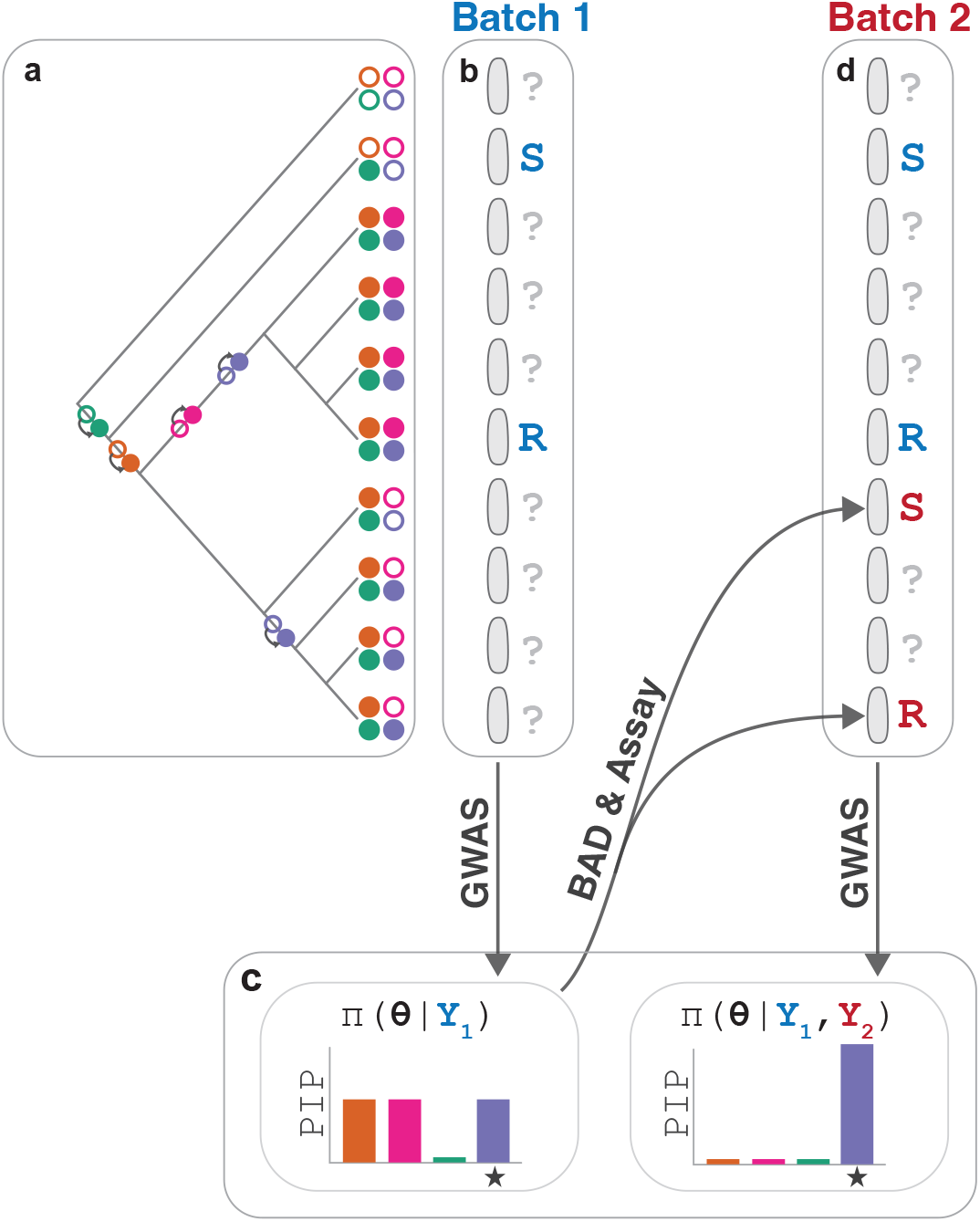
A conceptual diagram illustrating the application of BAD to GWAS for an antibiotic resistance phenotype. (a) The approach begins with a collection of sequenced isolates. Colored dots indicate the presence or absence of genetic variants superimposed on the phylogeny. (b) Assays to acquire phenotypic data are performed sequentially in batches. (c) The data from prior batches are used to update the posterior distribution of an underlying whole-genome regression model *π*(*θ* | *Y*_1_, …, *Y*_*n*_), with predictions from this model used to select the next batch of isolates to assay that maximally reduce the uncertainty in the model posterior, including the posterior inclusion probabilities (PIP). (d) The assay outcomes then inform iterative updates to the model until a candidate genetic variant reaches sufficient confidence to justify validation experiments, or until the maximum number of iterations is reached. The causal variant is indicated by a star.

### Uncertainty Quantification through a Bayesian Whole-Genome Regression Model

BAD requires a representation of uncertainty in the parameters that characterize genetic variants’ contribution to the phenotype of interest. To model phenotypic variation, we used a Bayesian sparse mixed model [26] adapted to binary phenotypes using a Probit likelihood [27]. We modeled phenotypic variation as a combination of large effects attributable to a small number of variants and the combined effect of genetic background. We first developed a version that employed a fixed scale for large effects, which we used in the experimental workflow to investigate *gyrB*^D429N^-mediated cross-resistance (Supplementary Text: Sparse Whole-Genome Regression Models). Because manually setting the scale of large effects can be challenging due to limited information, we later refined the model to estimate the scale. We used unitigs to represent genetic variation [28], although our approach is compatible with other representations, such as SNPs. Candidate variants are identified using posterior inclusion probability (PIP), which is the probability that at least one variant in a set of variants has a nonzero effect. PIPs can be computed for individual variants or for sets of variants, allowing aggregation by location [29] or by annotation.

We validated the whole-genome regression model using a surveillance dataset of *N. gonorrhoeae* [30] with three binary antimicrobial resistance phenotypes (cutoffs described in Figure 2) with well-characterized genetic determinants [31, 32] (Supplementary Text: Benchmark Phenotype Description). For each antibiotic, we sampled 800 isolates without replacement from the surveillance dataset to ensure a 10% prevalence of phenotypic resistance (Figure S1). Each dataset contained more than 100,000 variants with distinct occurrence patterns. When applied to these datasets, the model with a varying large effect scale successfully identified the genetic mechanisms of resistance (Figure 2, Supplementary Text: Regression Model Validation) across all scenarios. The results for the model with the fixed effect scale were practically indistinguishable (Figure S2). Our results outperformed the marginal linear mixed model (LMM) as implemented in pyseer [33] in precision and specificity across the three antibiotic datasets (Figure S3).

**Figure 2:**
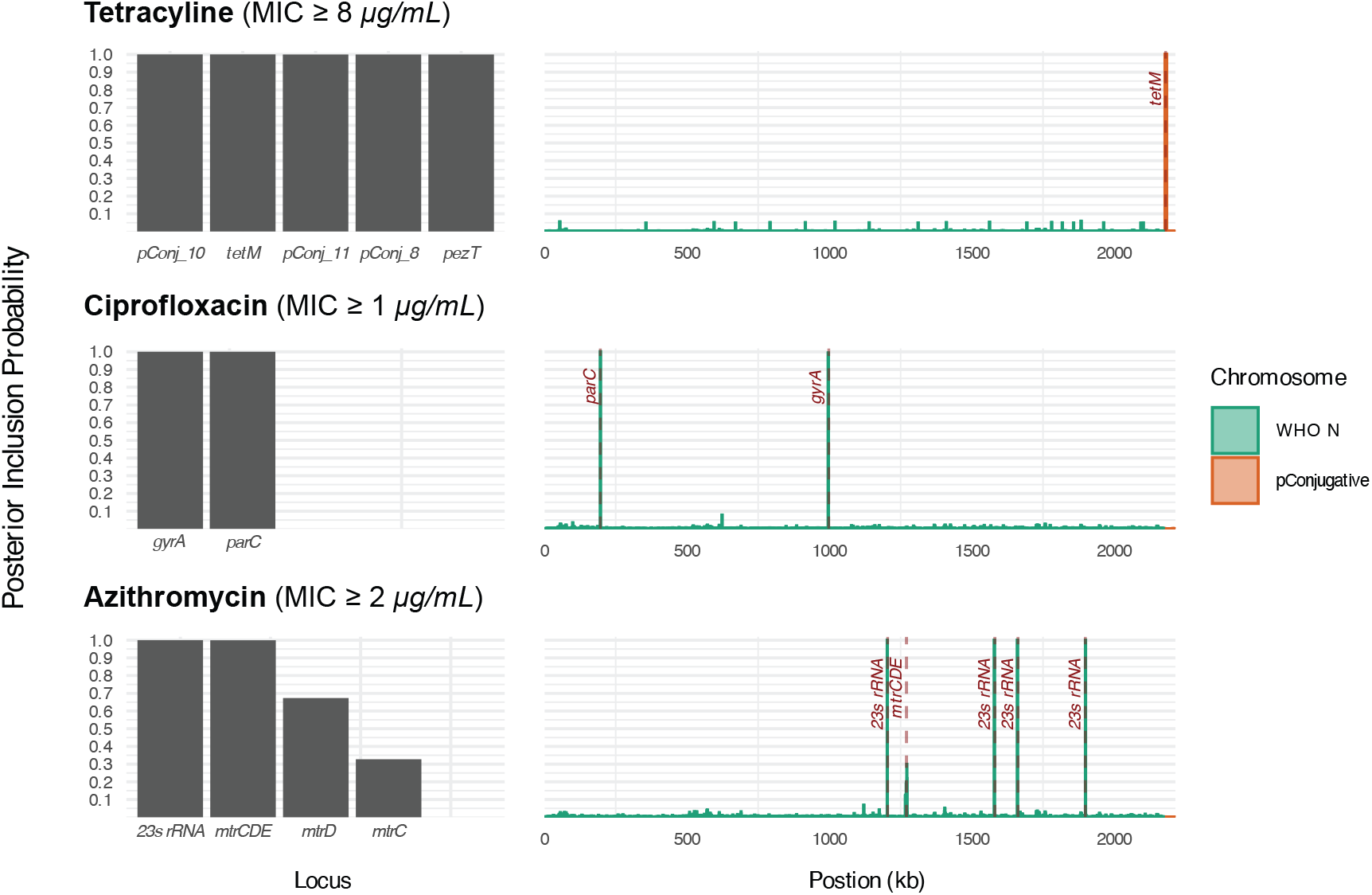
Validation of the Whole Genome Regression Model with Varying Large Effect Scale Using Three Antibiotic Resistance Phenotypes. Top: Tetracycline resistance. Middle: Ciprofloxacin resistance. Bottom: Azithromycin non-susceptibility. Left column: posterior inclusion probabilities (PIPs) for annotated genomic elements; shown: all annotated loci with PIP exceeding 0.25. Right column: PIPs for genomic regions, calculated for 1 kb overlapping windows spaced 500 bp apart. Locations of known causal loci are labeled using dashed lines and text. Shown: The WHO N chromosome and the conjugative plasmid. The gene *pezT* and the hypothetical proteins *pConj_8,10,11* flank the *tetM* insertion site and have perfect linkage with *tetM. mtrCDE* is the MtrCDE efflux pump operon.

### BAD enables highly efficient bacterial GWAS

We assessed the performance of BASS-GWAS in a series of synthetic experiments to identify variants associated with tetracycline resistance, ciprofloxacin resistance, or azithromycin non-susceptibility in *N. gonorrhoeae* using the same three datasets as before, with 20 simulated replicates for each experiment (Methods, Figure 2). We reasoned that prior evidence of phenotypic variation is a prerequisite for a successful bGWAS study. Therefore, for each replicate, we generated an initial seed batch of 8 strains by sampling one susceptible isolate and one resistant isolate, followed by randomly sampling six additional isolates from the remaining dataset without replacement. Starting from the initial set, in each replicate, we used either BAD or random sampling to acquire samples without replacement in batches of 8. We ran 9 iterations, yielding 72 additional isolates assayed per replicate.

BASS-GWAS with varying large-effect scale outperformed random sampling (Figure 3). For tetracycline resistance, all replicates using BAD identified the causal locus with PIP≥0.9 within 4 batches (Figures 3, S4), while only 30% of replicates using random sampling achieved PIP≥0.9 by batch 9 (Figures 3, S4). For ciprofloxacin resistance, all replicates using BAD identified both causal loci (*gyrA* and *parC*) with PIP≥0.9 by batch 9 (Figures 3, S5). In comparison, random sampling resulted in only 10% of replicates identifying *gyrA* and 25% of replicates identifying *parC* with PIP≥0.9 by batch 9 (Figures 3 and S5). For azithromycin non-susceptibility, the *mtrCDE* operon was identified with PIP≥0.9 by batch 9 in all replicates using BAD but in only 20% of the replicates using random sampling (Figures 3, S6). Neither BAD nor random sampling identified the 23S rRNA locus within 9 batches (corresponding PIP < 0.1 at all times) (Figure 3), likely because of its low prevalence in our dataset: only 15/800 isolates (<2%) carried two or more copies of 23S rRNA gene with the C2611T substitution [34]. The superior performance of BASS-GWAS sampling was consistent across a range of PIP cutoffs for each of the antibiotics (Figure 3, Figures S5,S4, S6), demonstrating the efficiency gains attributable to BAD. The performance of BASS-GWAS with a fixed large-effect scale was similar to its performance with varying large-effect scale, with the exception that the fixed large-effect scale yielded oscillatory behavior for the PIPs of *gyrA* and *parC* in the ciprofloxacin benchmark (Figures S7, S8).

**Figure 3:**
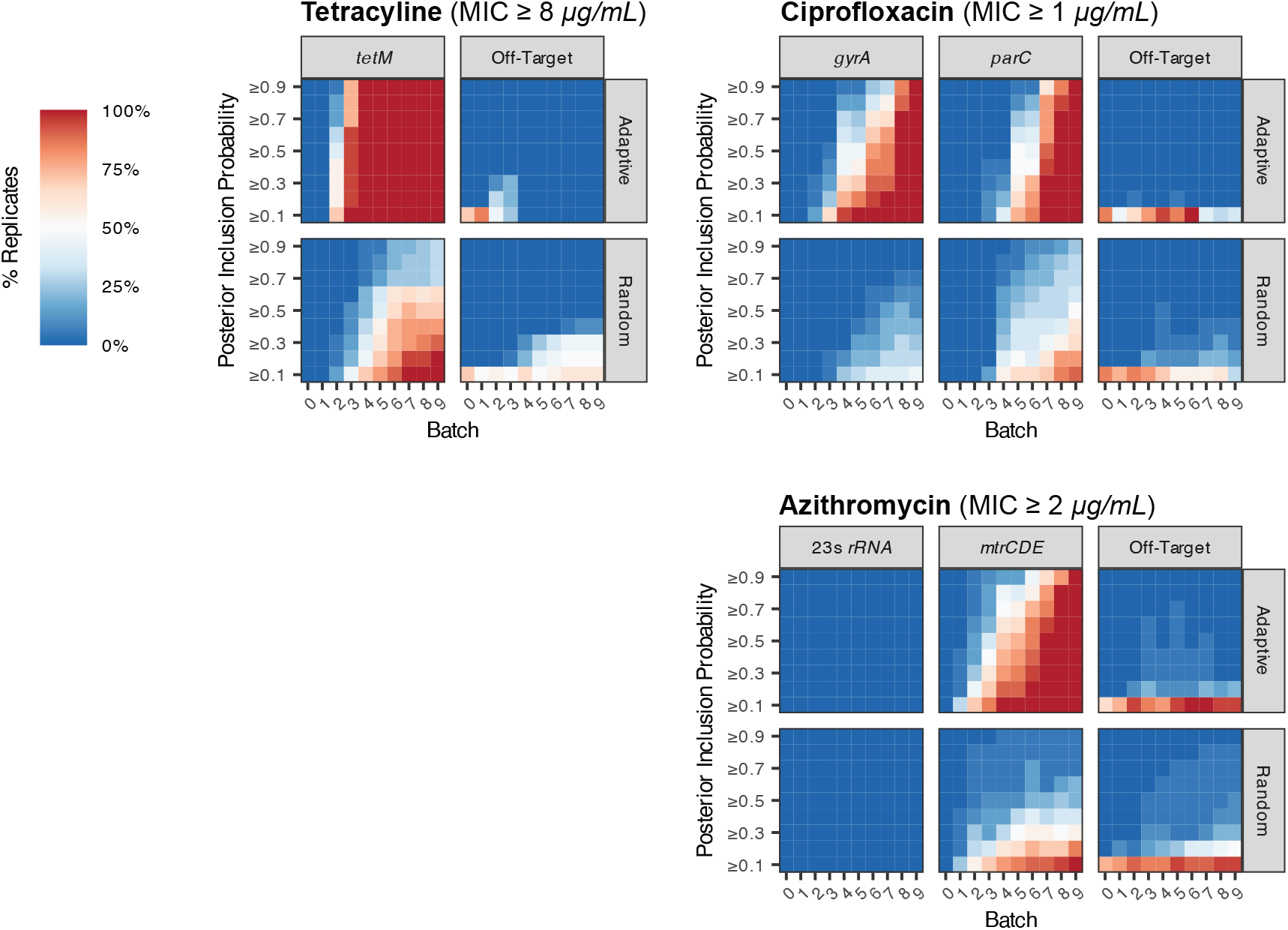
Comparison of Adaptive and Random Sampling. The heatmap depicts the percentage of replicates (20 simulated experiments) for which the PIP of an annotated gene exceeded each threshold after assaying a given number of batches (8 strains per batch), highlighting the PIPs of known causal loci and the highest PIP among all remaining annotations (denoted Off-Target). Causal locus for tetracycline resistance: *tetM*. Causal loci for ciprofloxacin resistance: *gyrA* and *parC*. Causal loci for azithromycin non-susceptibility: the *mtrCDE* operon and 23S rRNA. Batch 0 consists of the initial randomly selected samples.

We evaluated the effect of batch size on BAD efficiency using the ciprofloxacin resistance dataset. We compared the experiments described above, which used a batch size of 8, to experiments with batch sizes of 4 and 16. We applied BASS-GWAS with varying large effect scale. Adaptive sampling outperformed random sampling for all tested batch sizes (Figure S9). For batch sizes of 4 and 8, the number of strains required to identify the *gyrA* locus in at least 70% of the replicates using a PIP threshold of 0.9 was 68 and 64, respectively (Figure S10). For *parC*, the number of additional strains required was 64 and 64, respectively (Figure S10). However, a batch size of 16 reduced the efficiency of adaptive sampling (Figure S10): *gyrA* and *parC* loci were identified in at least 70% of the replicates using a PIP threshold of 0.9 only after 80 strains (Figure S10). This reduction in efficiency was consistent with theoretical expectations (Supplementary Text: Impact of Batch Size). We did not assess batch sizes larger than 16, but we verified that batch sizes of up to 20 were computationally tractable.

To assess how BASS-GWAS performs in highly imbalanced scenarios, we generated an additional tetracycline resistance dataset of 800 isolates in which only 2.5% were resistant (Methods, Figure S1) and used BASS-GWAS with varying large effect scale. To establish a baseline, we first applied the whole-genome regression model to this dataset with phenotypic information for all 800 isolates. The model correctly identified the causal *tetM* locus with high probability (Figure S11). BASS-GWAS replicates with batch size 8 identified the causal locus with PIP≥0.9 by batch 4 (Figures S12, S4). None of the replicates using random sampling identified the causal locus with PIP≥0.9 by batch 9 (Figures S12, S4).

### *parC*D86N enables *gyrB*D429N-mediated gepotidacin resistance

Having demonstrated the effectiveness of BASS-GWAS on synthetic datasets, we next applied BASS-GWAS to identify the genetic variants underlying *gyrB*^D429N^-mediated gepotidacin resistance in *N. gonorrhoeae*. For consistency, we used the model with fixed large effect scale for all batches. The seed dataset comprised nine diverse clinical isolates previously tested [25], two of which demonstrated cross-resistance (>4-fold increase in gepotidacin MIC following the introduction of *gyrB*^D429N^). We performed BASS-GWAS with a test batch size of 4 strains (Figure 4a,b, Supplementary Text: BASS-GWAS for the genetic basis of zoliflodacin/gepotidacin cross-resistance), with one exception: in the first iteration, we used BAD to select 3 strains and manually pre-selected the fourth strain to represent a clade with an unusual *gyrA* allele, *gyrA*^S91F/A92P/D95Y^. This *gyrA* allele contains a proline substitution at position 92, a key residue modulating gepotidacin susceptibility [24]. Isolates from this *gyrA*^S91F/A92P/D95Y^ clade have increased resistance to both sitafloxacin [35] and gepotidacin [36]. Because strains from this clade were not represented in the seed dataset and were present in only a small fraction of strains in our laboratory collection, we manually selected strains to ensure this genotype of concern was included in the experiment.

**Figure 4:**
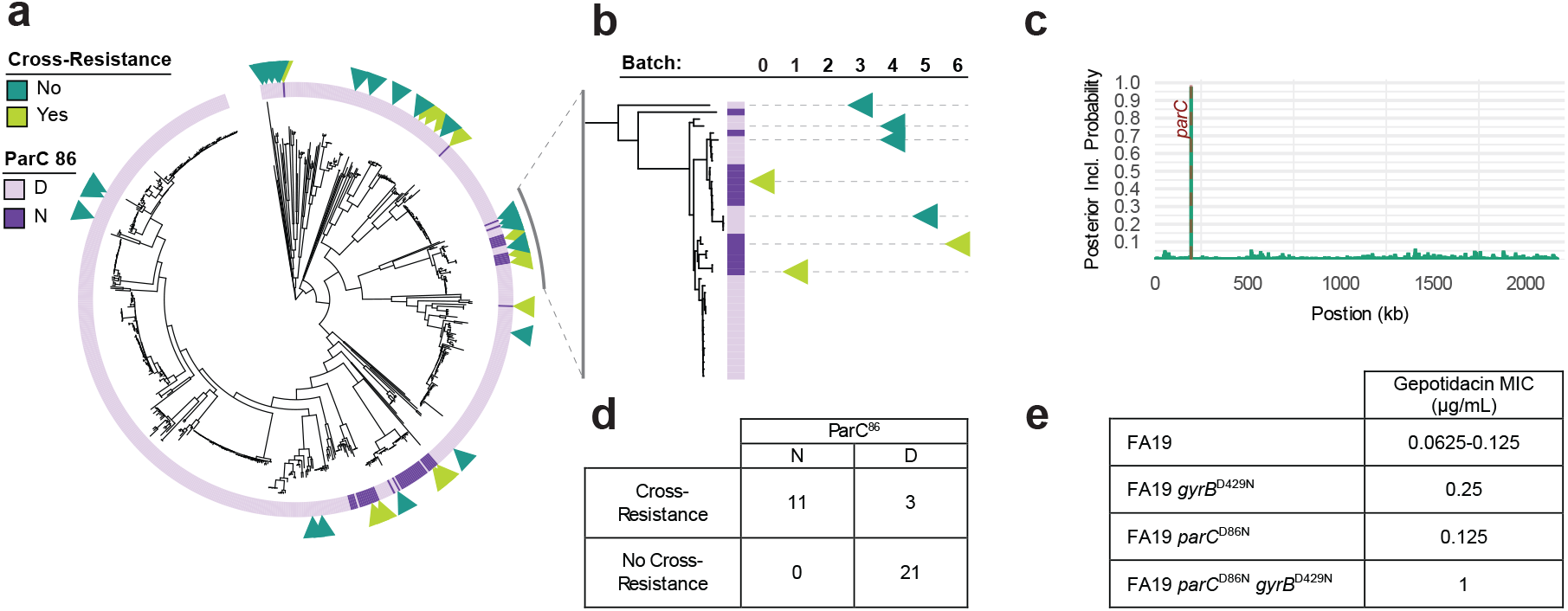
BASS-GWAS identification of *parC*^D86N^ as a candidate variant enabling *gyrB*^D429N^-mediated gepotidacin resistance. (**a**) Tree of strains available to the GWAS algorithm. Outer annotation: results of *gyrB*^D429N^-mediated gepotidacin resistance phenotyping assays from the seed dataset and 26 additional tested strains (triangles, colored by phenotype). Inner ring: *parC* ^86^ allele, with the *parC* ^86N^ variant associated with cross-resistance shown in dark purple. (**b**) Example of adaptive sampling in one highly sampled clade over the course of the experiment. Triangles represent phenotype information for strains present in the seed dataset (batch 0) or chosen in subsequent experimental batches. (**c**) Manhattan plot of the posterior inclusion probability for each 1 kb genomic region following completion of the GWAS experiments (7 batches). The genomic region corresponding to *parC* (PIP 0.969 for the 1 kb region) is labeled in red. Shown: the WHO N chromosome. (**d**) Phenotypic results of all tested strains from the seed dataset and the 7 adaptively selected batches, stratified by *parC* ^86^ allele. (**e**) Range of gepotidacin MICs of *parC* and *gyrB* mutants in the laboratory strain FA19 from two independent experiments.

After batch 7, BASS-GWAS identified an association between *gyrB*^D429N^-mediated gepotidacin resistance and a single variant encoding *parC*^D86N^ (PIP=0.953) (Figures 4c, S13, S14). All tested strains with this variant demonstrated cross-resistance (n=11, Figure 4, table S1), and the presence or absence of this variant correlated perfectly with the presence or absence of cross-resistance in the seed dataset. Cross-resistance was also observed in 3 isolates that lacked *parC*^D86N^ (Figure 4d).

To test whether *parC*^D86N^ is sufficient to enable zoliflodacin/gepotidacin cross-resistance, we introduced *parC*^D86N^ into the *N. gonorrhoeae* laboratory strain FA19. As expected, there was no effect of *parC*^D86N^ on the gepotidacin MIC of FA19 in the absence of *gyrB*^D429N^ (Figure 4e), and introduction of the zoliflodacin resistance mutation *gyrB*^D429N^ alone also did not substantially increase gepotidacin MICs in the wildtype FA19 strain (<4-fold increase). However, introduction of *gyrB*^D429N^ into the FA19 *parC*^D86N^ strain increased the gepotidacin MIC 16-to 32-fold (Figure 4e).

### *gyrBM*^D429N^-mediated gepotidacin resistance in a clade lacking *parC*^D86N^

We observed *gyrB*^D429N^-mediated gepotidacin resistance in 3 strains that did not carry the *parC*^D86N^ variant: HHH012, EEE018, and GGG005. These strains belong to the only clade in our collection that carried the *gyrA*^S91F/A92P/D95Y^ allele. These strains have unusually high gepotidacin MICs at baseline (1-2 *μ*g/mL) with ≤8-fold increases in gepotidacin MICs following the introduction of *gyrB*^D429N^ (table S1). BASS-GWAS did not identify any variants with high confidence as candidate drivers of this lineage-specific phenotype.

Because *gyrA*^A92T^ is implicated in gepotidacin resistance [24], we hypothesized that the *gyrA*^S91F/A92P/D95Y^ variant might contribute to gepotidacin resistance in this lineage, either alone or in epistasis with *gyrB*^D429N^. The combination of *gyrA*^S91F/A92P/D95Y^ and *gyrB*^D429N^ selected for off-target mutations in FA19, so we instead tested the effects of these alleles in GCGS0481, a strain that tolerates diverse *gyrA* alleles and *gyrB* mutations [37, 38]. In GCGS0481, the *gyrA*^S91F/A92P/D95Y^ allele did not directly increase gepotidacin MIC at baseline, nor was it sufficient to enable *gyrB*^D429N^-mediated gepotidacin resistance (table 1).

**Table 1:** *gyrA*^S91F/A92P/D95Y^ and *parE*^D437N^ contribute to gepotidacin resistance. Gepotidacin MICs of *gyrA, gyrB*, and *parE* mutants in a ciprofloxacin-susceptible derivative of clinical strain GCGS0481 and in the clinical strain EEE018 (* denotes that strain contains a kanamycin marker upstream of *gyrA*; ** denotes that strain contains a kanamycin marker downstream of *parE*). Bolded alleles have been changed from the parental genotype.

| Strain background | GyrA<sup>91/92/95</sup> | GyrB<sup>429</sup> | ParE<sup>437</sup> | Gepotidacin MIC ($\mu\text{g/mL}$) |
| --- | --- | --- | --- | --- |
| GCGS0481 *gyrA*<sup>S/D</sup> | S/A/D | D | D | 0.125-0.25 |
| GCGS0481 *gyrA*<sup>S/D</sup> | S/A/D | **N** | D | 0.125-0.25 |
| GCGS0481 *gyrA*<sup>S/D</sup> | **F/P/Y** | D | D | 0.125-0.25 |
| GCGS0481 *gyrA*<sup>S/D</sup> | **F/P/Y** | **N** | D | 0.25 |
| EEE018* | F/P/Y | D | N | 2-4 |
| EEE018* | F/P/Y | **N** | N | 16-32 |
| EEE018* | **S/A/D** | D | N | 1-2 |
| EEE018* | **S/A/D** | **N** | N | 4-8 |
| GCGS0481 *gyrA*<sup>S/D</sup> | S/A/D | D | **N** | 1 |
| GCGS0481 *gyrA*<sup>S/D</sup> | S/A/D | **N** | **N** | 2 |
| GCGS0481 *gyrA*<sup>S/D</sup> | **F/P/Y** | D | **N** | 1-2 |
| GCGS0481 *gyrA*<sup>S/D</sup> | **F/P/Y** | **N** | **N** | 2 |
| EEE018** | F/P/Y | D | N | 2 |
| EEE018** | F/P/Y | **N** | N | 16 |
| EEE018** | F/P/Y | D | **D** | 2 |
| EEE018** | F/P/Y | **N** | **D** | 4-8 |

We then tested whether *gyrA*^S91F/A92P/D95Y^ is necessary for *gyrB*^D429N^-dependent gepotidacin cross-resistance in its native genomic context. We replaced *gyrA*^91F/92P/95Y^ with the wild-type *gyrA*^91S/92A/95D^ allele in EEE018, one of the previously tested strain from this lineage. *gyrA*^91S/92A/95D^ reduced the the baseline gepotidacin MIC of EEE018 by 2-fold. Subsequent introduction of *gyrB*^D429N^ into EEE018 *gyrA*^91S/92A/95D^ then increased the gepotidacin MIC 4-fold, compared to an 8-fold increase in parental EEE018 (table 1). This partial reversal of gepotidacin/zoliflodacin cross-resistance suggests that cross-resistance in this lineage is a polygenic trait to which *gyrA*^S91F/A92P/D95Y^ contributes. We therefore sought to identify other contributing variants.

Outside the *gyrA*^91F-95Y^ region, EEE018 had no candidate variants in *gyrA, gyrB*, and *parC*, as all nonsynonymous variants in these loci were also found in strains that lacked cross-resistance. However, all strains in the EEE018 clade share a *parE*^D437N^ substitution. ParE is a homolog of GyrB that complexes with ParC to form topoisomerase IV. Moreover, *parE*^437^ is homologous to *gyrB*^429^. To determine whether *parE*^D437N^ affects gepotidacin resistance, we introduced *parE*^D437N^ into the GCGS0481 background with and without *gyrA*^S91F/A92P/D95Y^. The *parE*^D437N^ allele increased the baseline gepotidacin MIC 4-to 8-fold. However, *parE*^D437N^ did not enable *gyrB*^D429N^-mediated gepotidacin cross-resistance, with at most a 2-fold additional increase in gepotidacin MIC resulting from introduction of *gyrB*^D429N^ into these strains (table 1).

To investigate the contribution of *parE*^D437N^ in its native genetic background, we replaced it with the wild type *parE*^437D^ allele in EEE018. In contrast to the results in GCGS0481, the baseline gepotidacin MIC of EEE018 *parE*^437D^ was unaltered. However, introduction of *gyrB*^D429N^ into the EEE018 *parE*^437D^ strain resulted in only a 2-to 4-fold increase in gepotidacin MIC, compared to 8-fold in parental EEE018 (table 1).

## Discussion

Bacterial GWAS has uncovered the genetic mechanisms underlying a range of resistance and virulence phenotypes [1, 4, 5, 7]. While genome sequencing has historically accounted for much of the cost and effort involved in bacterial GWAS, in recent years, large collections of sequenced strains have become commonplace. These collections can now enable efficient discovery of the genetic basis of phenotypes that have been more challenging to test.

To achieve this efficiency, we developed BASS-GWAS, combining BAD with a whole-genome regression model. Validation across a range of real-world antimicrobial resistance phenotypes in *N. gonorrhoeae* demonstrated that the whole-genome regression model enables precise genotype-to-phenotype mapping and that combining this model with BAD allows for highly sample-efficient bacterial GWAS. BASS-GWAS vastly outperformed random sampling across resistance phenotypes that differed in penetrance and genetic basis, greatly reducing the number of samples required to correctly identify causative genetic variants. For example, when applied to ciprofloxacin resistance, all simulated replicates using adaptive sampling identified both causal loci – *gyrA* and *parC* – with PIP>0.9 after analysis of 80 isolates. In contrast, simulated random sampling of 160 isolates identified *parC* with a PIP>0.9 in 60% of replicates and *gyrA* in only 35%.

We used BASS-GWAS to investigate which variants underlie strain-dependent *gyrB*^D429N^-mediated cross-resistance between zoliflodacin and gepotidacin [25]. Taking advantage of the flexibility offered by BASS-GWAS, we conditioned the initial design to include a pre-selected isolate from a clade of concern. After 7 batches of isolates, representing 26 additional strains, we identified *parC*^D86N^ as associated with cross-resistance. We experimentally validated that *parC*^D86N^ is sufficient to confer *gyrB*^D429N^-mediated cross-resistance. Moreover, we identified another pathway to cross-resistance within a single clade of isolates without the *parC*^D86N^ variant. This clade had no closely related phylogenetic neighbors in our isolate collection, and low homoplasy precluded the successful application of further GWAS. Hypothesis-driven experimental genetics allowed us to characterize cross-resistance in this lineage as a polygenic phenotype arising from *gyrA*^S91F/A92P/D95Y^, *parE*^D437N^, and possibly other variants.

While more work remains to fully characterize these genetic pathways, our findings underscore the complexity of possible pathways to resistance to topoisomerase-targeting antibiotics in *N. gonorrhoeae*. Decades of fluoroquinolone use have left a legacy of extensive genetic variation in topoisomerase components, with incompletely understood impacts on strain fitness and resistance profiles in an evolutionary landscape of shifting antimicrobial pressures. As additional strain backgrounds are studied, we may discover additional pathways to gepotidacin and zoliflodacin resistance.

There are limitations to our approach. The first is the dichotomization of continuous phenotypes. Using a binary likelihood enables highly sample-efficient EIG estimators [39], simplifying experimental-design optimization. However, while many phenotypes of interest are binary or can be reasonably dichotomized, dichotomizing truly continuous phenotypes reduces statistical efficiency. The second limitation is the batch size. This follows from the EIG estimator’s exponential complexity with respect to batch size. Several emerging strategies aim to improve the efficiency of EIG estimation for continuous likelihoods or large batch sizes [16], thereby facilitating the development of more flexible BASS-GWAS.

As collections of microbial isolates with genome sequences grow, the experimental effort behind microbial GWAS will shift from sequencing to characterizing phenotypes that are increasingly challenging to assay, emphasizing the importance of efficient experimental design. Here, we achieve this goal by introducing BASS-GWAS, an approach based on BAD. By iteratively selecting strains to phenotype, BASS-GWAS incorporates information from intermediate experiments and enables bacterial GWAS with as few as tens of isolates, which is critical when studying hard-to-measure phenotypes.

## Methods

### Benchmark Dataset Construction

To construct the four benchmark datasets, we used publicly available genomes and associated metadata [30]; see (table S2).

### Genomic Analysis

*De novo* assembly was performed using SPAdes version 3.12.0 [40] with the `--careful` flag, and reference-based mapping to NCCP11945 (NC_011035.1) was done using BWA-MEM version 0.7.17 [41]. We used Pilon version 1.23 to call variants (minimum mapping quality: 20, minimum coverage: 10X) [42] after marking duplicate reads with Picard version 2.20.1 (https://broadinstitute.github.io/picard/) and sorting reads with samtools version 1.17 [43]. We generated pseudogenomes by incorporating variants supported by at least 90% of reads and sites with ambiguous alleles into the reference genome sequence. We mapped reads to a single copy of the locus encoding the 23S rRNA and called variants using the same procedures [34]. *De novo* assemblies were annotated using Prokka version 1.14.6 [44].

We identified resistance-associated alleles from *de novo* assemblies and pseudogenomes. We identified single-nucleotide variants (e.g., mutations in *gyrA, parC, ponA*, and *penA*) and the copy number of resistance-associated variants in 23S rRNA from variant calls. To determine the presence or absence of genes, mosaic alleles, promoter variants, and small insertions or deletions, we used the results of blastn version 2.9.0 [45] searches of assemblies for resistance-associated genes.

To construct recombination filtered phylogenies, we used Gubbins version 3.3.1 [46] and IQTree version 2.4.0 [47].

### GWAS Input

We used unitigs [28] to represent genetic variation. To generate unitigs from *de novo* assemblies, we used unitig-caller, which is part of pyseer [33]. To obtain the input matrix *X* from the binary matrix that encoded the presence and absence of unitigs, we: (1) complemented the presence patterns of any unitigs that were present in more than 50% of all samples; followed by (2) collapsing unitigs with identical presence-absence patterns into a single variant; and (3) centering each column by subtracting its mean from each entry. Each variant corresponded to potentially multiple unitigs with the same presence-absence pattern. To represent the population structure, we used a low-rank approximation *C* to the phylogenetic correlation matrix obtained by truncating its singular value decomposition to the top-*k* singular values that account for 99% of the variance. We annotated and mapped all unitigs to reference genomes using pyseer [33].

To associate each variant with genomic location, we mapped unitigs to the WHO N reference genome [48]. To associate each variant with genomic annotations, we mapped unitigs to the WHO N, WHO G, WHO M, and WHO F reference genomes [48] and the *de novo* assemblies of the genomes that comprised the respective datasets. In both cases, we used pyseer utilities to map unitigs [33].

### Whole Genome Regression Model

We used an *N* ×*M* matrix *X* to encode the presence and absence of *M* genetic variants in a collection of *N* isolates. The *N* × *N* matrix *C* encoded the population structure. We specified observations (*y*_*i*_, *ξ*_*i*_) by the index of the assayed strain, *ξ*_*i*_, and the observed phenotype, *y*_*i*_. The model is given by equation 1:

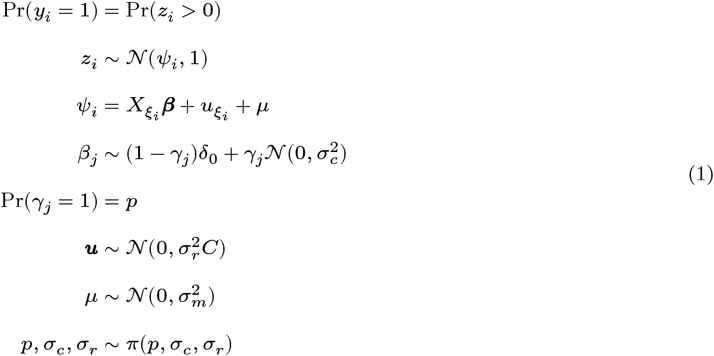

Here, 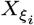 denotes the row of *X* indexed by *ξ*_*i*_, *β* represents the effect of genetic variants, and lineage effects are represented by 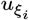 for each strain 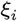. For each variant *j, γ*_*j*_ is an indicator variable equal to 1 if *β*_*j*_ has a non-zero effect. The indicators *γ* are independently and identically distributed, with the probability that *γ*_*j*_ = 1 equal to *p*, which governs the proportion of large effects. The proportion of large effects *p* follows a beta prior. To model non-zero large genetic effect sizes, we used a normal distribution with variance 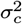. To model lineage effects, we used a normal distribution with variance 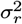.

We initially generated a version of this model with a fixed scale for large effects, in which *σ*_*c*_ was specified rather than fit. This is a common assumption in the Bayesian variable selection literature [49]. In this fixed-scale model, hyperparameters were defined as follows:

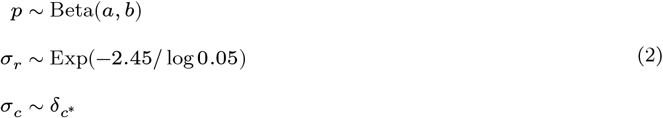

To define prior parameters *a* and *b* for the distribution *p* governing the proportion of variants with large effects, we used the *a priori* mean proportion of causal variants, *μ*_*p*_, and a pseudo-dispersion, *ϕ*_*p*_: *a* = *μ*_*p*_*ϕ*_*p*_ and *b* = (1 −*μ*_*p*_)*ϕ*_*p*_. We used *μ*_*p*_ = 5/*M*, reasoning that 5 represents a conservative choice for the average prior number of causal variants. We used *ϕ*_*p*_ = *M*/10 to penalize overfitted models. For *σ*_*r*_, which represents the variance of lineage effect sizes, we used a penalized complexity prior [50] for 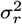 parameterized to constrain approximately 95% of the prior mass for 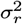 to be less than 2.45^2^ ≈ 6. To define *σ*_*c*_, which represents the variance of effect sizes for genetic variants with large effects, we set the hyperparameter 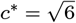. This corresponds to a prior probability of approximately 10% of observing a variant with an absolute effect size exceeding 4. An effect size of 4 has been argued to be reasonable for high-penetrance resistance-conferring variants [51]. We set 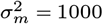 as a non-informative choice for the intercept term *μ*.

As a fixed 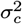 can lead to model misspecification, and the scale of large effects may vary between organisms and phenotypes, we adapted our model to estimate the scale of large effects and reparameterized the prior for *p*:

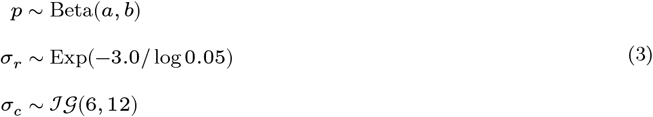

Here, we parameterized *p* in terms of the *a priori* probability *p*_0_ that there are 0 large effects, and the *a priori* average model size *m*_*p*_, conditional on there being at least one effect. To find the values of *a* and *b* that satisfy *p*_0_ and *m*_*p*_, we used numerical optimization. We used *p*_0_ = 0.25, reasoning that most studies will have at least one causative variant. We set *m*_*p*_ = 5, reasoning that 5 represents a conservative choice for the average prior number of causal variants. As above, to estimate lineage effect sizes we used a penalized complexity prior [50] for 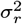, in this case constraining approximately 95% of the prior mass for 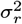 to be less than 9. We equip the variant effect scale parameter *σ*_*c*_ with an inverse gamma distribution *JG*(6, 12). We constrained the prior variance for the intercept parameter *μ* to 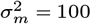 to avoid placing excessive prior mass on scenarios with no phenotypic variation.

The model structure yields the following posterior factorization:

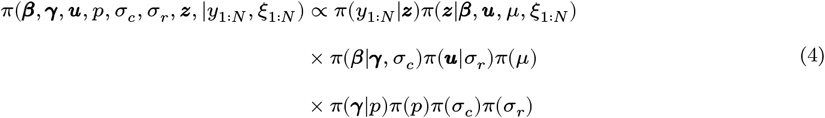

Further, by marginalizing *β, u, μ, p* we obtained:

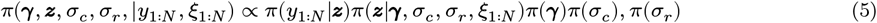

We implemented posterior sampling based on the weighted Tempered Gibbs Sampling (wTGS) [52], incorporating modifications [53, 54]. Briefly, for a posterior distribution *π*(*θ*|*y*_1:*N*_, *ξ*_1:*N*_) wTGS targets a *tempered* distribution *f*(*θ*) = *Z*(*θ*)*π*(*θ*|*y*_1:*N*_, *ξ*_1:*N*_) that is constructed so that its geometry makes it easier to sample from, while remaining close to the untempered target *π*(*θ*|*y*_1:*N*_, *ξ*_1:*N*_). Expectations with respect to the original target can be computed by using self-normalized importance sampling (SNIS) [52]. See (Supplementary Text: MCMC Sampling) for a detailed description of the sampling scheme.

### Posterior summaries of genetic associations

We considered three posterior summaries to evaluate genetic associations. The first summary was the posterior inclusion probability (PIP) of each variant, defined as the posterior probability that a variant has a non-zero effect on the measured phenotype. The PIPs of variants represent the most granular level of mapping, directly assigning a probability that a given variant is causal. A downside of this granularity is that variant PIPs can be difficult to interpret when there is substantial uncertainty about which specific variant is causal, as is often the case with unitigs. The second summary we considered was the PIPs of genomic regions, which aggregate the effect of nearby variants and represent the posterior probability that a given genomic region of a set size contains at least one variant with a non-zero effect [29]. The PIPs of genomic regions are highly interpretable and offer control over granularity by adjusting the window size. The downside of the PIPs of genomic regions is reference bias: highly divergent variants will not map to a single reference. Third, we considered the PIPs of annotated regions, which aggregate variants mapping to loci with the same annotation across multiple reference genomes and represent the posterior probability that a genomic region with a given annotation contains at least one variant with a non-zero effect.

The PIP of a variant *V* is defined as the following posterior probability:

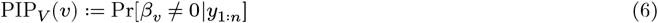

To estimate PIP_*v*_ (*V*) we used a Rao-Blackwellized Monte-Carlo (MC) estimate that is computed as a part of the MCMC sampler used [52].

To define the PIP of a genomic region *g*, we first introduce the set of all variants that contain at least one unitig overlapping with a genomic region *g, S*_*G*_(*g*). The PIP of a genomic region *g* is then defined as the following posterior probability:

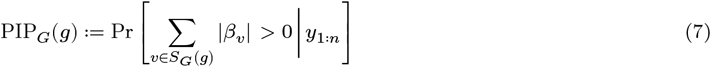

To determine *S*_*G*_(*g*), we used the mapping of unitigs to the WHO N reference genome [48].

To define the PIP of a genomic annotation *a*, we first introduce the set of all variants that contain at least one unitig annotated with *a*, denoted *S*_*A*_(*a*). The PIP of a genomic annotation *a* is then defined as the following posterior probability:

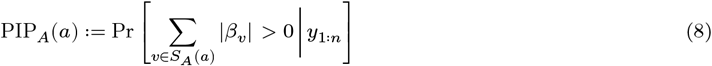

For both PIP_*G*_ and PIP_*A*_, we used standard MC estimators. To determine *S*_*A*_(*a*), we used the unitig annotations.

### Bayesian Adaptive Design - EIG Estimation

BAD requires evaluating the incremental EIG to optimize experimental designs. Given an experimental history *h*_*n*_ := (*y*_1:*N*_, *ξ*_1:*n*_) of strain choices *ξ*_1:*n*_ = *ξ*_1_, *ξ*_2_, …, *ξ*_*n*−1_, *ξ*_*n*_ and the associated experimental outcomes *y*_1:*N*_ = *y*_1_, *y*_2_, …, *y*_*n*−1_, *y*_*n*_, the incremental EIG of a batch of *k* unobserved strain indices *ξ*_*n*+1:*n*+*k*_ = *ξ*_*n*+1_, *ξ*_*n*+2_, …, *ξ*_*n*+*k*−1_, *ξ*_*n*+*k*_ is defined as [14, 16]:

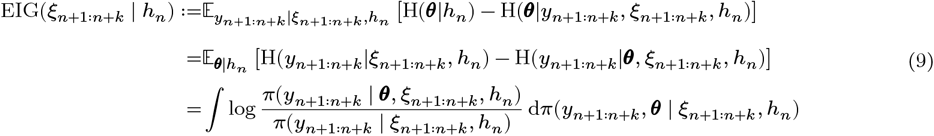

Here, H(·) denotes the entropy. As the EIG involves a doubly-intractable integral, it must be estimated. We aimed to estimate the EIG with respect to the full set of model parameters: (*γ, β, u, μ, σ*_*r*_, *σ*_*c*_). As a consequence of the conditional independence between *y* and (*γ, σ*_*r*_, *σ*_*c*_) given (*β, u, μ*), this is equivalent to estimating the EIG with respect to *θ* = (*β, u, μ*). We used a Rao-Blackwellized estimator applicable to models with discrete observations [39, 55]. Given draws *θ*^(1:*M*)^, *Z*^(1:*M*)^ ~ *Z*(*θ*)*π*(*θ* | *h*_*n*_) obtained using wTGS sampling, the estimator is given by (Equation 10).

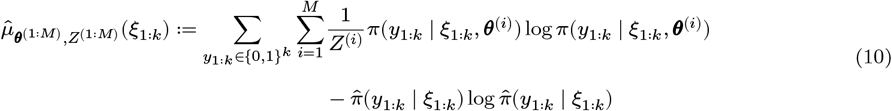

Where

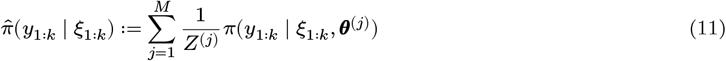

The structure of the estimator is analogous to that of the estimator used in BatchBALD [55]. We used the same evaluation strategy as in BatchBALD, storing intermediate likelihood matrices to speed up computations.

### Bayesian Adaptive Design - Batch Optimization

At each step, we sought to identify a batch 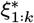 of *k* distinct isolates for the next round of assays to maximize the incremental EIG (Equation 9). This myopic maximization of the EIG is a common approach in applications of BAD [16].

Optimizing a batch of isolates to maximize the EIG exactly is a combinatorial optimization problem and therefore computationally intractable. However, for models with conditionally independent observations, the EIG is a monotone and submodular function [55] (Supplementary Text: 1 − 1/*e* Optimality of EIG Optimization). This allows for an approximate 1−1/*e* optimal solution to be obtained using greedy maximization [55, 56] (Supplementary Text: 1 − 1/*e* Optimality of EIG Optimization). We denoted the set of *N* strains in the collection, both observed and unobserved, as *S* = {1, 2, .., *N* − 1, *N*} and the set of unobserved strains by *S*_*r*_ ⊂ *S*_*o*_. We aimed to select a set of *k* distinct isolates *ξ*_1:*k*_ from *S*_*r*_ for the next batch of experiments. We set *ξ*_0_ = ∅ and iteratively choose isolates *ξ*_1_, *ξ*_2_, …, *ξ*_*k*−1_, *ξ*_*k*_ to maximize the increment of the EIG estimate:

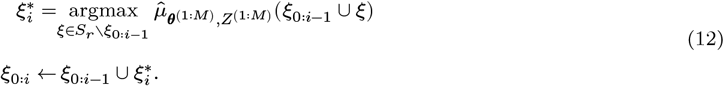

### Implementation

We implemented BASS-GWAS as a julia [57] package. The package, together with helper scripts implemented in R [58].

### Whole Genome Regression Model Validation

We ran both versions of BASS-GWAS on the respective dataset, providing phenotypic information for all 800 samples, with the experimental design turned off. We configured posterior sampling with 6 parallel chains, each running 750×1000 iterations and sampling every 750 iterations. This yielded 6000 posterior samples in total.

### Marginal LMM Comparison

We ran pyseer LMM [33] on each of the three benchmark datasets. We filtered significantly associated variants with a Bonferroni-corrected p-value < 0.05. We then plotted the resulting significantly associated variants as a Manhattan plot.

### Experimental Design Benchmarks

To evaluate the performance of BASS-GWAS, we conducted a series of simulation experiments. Each experiment consisted of 20 replicates. When using a batch size *k* we generated an initial set of *k* strains for each replicate by first sampling one susceptible isolate and one resistant isolate at random. Next, we sampled *k*− 2 additional isolates from the remaining data set without replacement. Starting from the initial set of strains, in each replicate we: (1) used BAD to acquire samples in batches of k, and (2) used random sampling to acquire samples in batches of k. In both cases, we acquired samples without replacement. For each replicate, we configured posterior sampling to use 4 parallel chains, each for 500×1000 iterations, sampling every 500 iterations. This yielded 4000 posterior samples in total. The typical run time per iteration per replicate was between 2 and 4 hours.

We first conducted one experiment for each of the 3 resistance datasets using a batch size of *k* = 8. To further evaluate the impact of varying batch size, we conducted two additional experiments using the ciprofloxacin resistance dataset, each with a batch size of *k* = 4 or *k* = 16.

When using batch sizes of *k* = 8 and *k* = 16, we ran each experiment for 9 iterations, yielding 72 and 144 additional isolates, respectively, per replicate and sampling scheme. With a batch size of k=4, we ran each experiment for 19 iterations, yielding 76 additional isolates per replicate and sampling scheme.

### Iterative Experimental Design

We compiled sequencing data for (n=735) isolates in our strain collection. We used BASS-GWAS to select isolates in batches of 4. To obtain posterior draws, we ran 8 chains in parallel, each for 750×1500 iterations, sampling every 750 iterations. The combined runtime for sampling and experimental design was under 8 hours. For the first batch, we manually fixed one isolate and then used BAD to select the remaining three, conditional on the first isolate. After optimizing the second batch, we noticed that some of the selected strains were missing or mislabeled. Therefore, we conducted a strain inventory audit, which identified 11 missing or mislabeled strains. We re-generated the GWAS input using the remaining (n=724) strains, and optimized a new batch. During preparation of batch 5, we found that the frozen stock and genome sequence for one selected strain resulted from a mixed culture of multiple *N. gonorrhoeae* strains. We therefore ran additional quality checks on the genome assemblies of each strain in our collection and removed isolates with ambiguous SNP calls at resistance-linked loci. This removed 26 additional isolates.

### *N. gonorrhoeae* Strains, Culture Conditions, and MIC Measurement

All strains are described in (table S1). Strains were grown on GCB agar (Difco) supplemented with Kellogg’s supplement [59] (GCB-K) at 37°C with 5% CO_2_. Gepotidacin and zoliflodacin MICs were measured via agar dilution in GCB-K (two replicates per experiment).

### Primers

All primers used are described in (table S4).

### Introduction of *gyrB*^D429N^ into experimental strains for BASS-GWAS

The *gyrB*^D429N^ allele was isolated from YG0050, which acquired the *gyrB*^D429N^ allele during experimental evolution under ciprofloxacin pressure (described as Experimental evolution GCGS0481_SG isolate 3 in [37]) via PCR with primers AM_1 and AM_2. Strains were transformed with approximately 500 ng of the resulting *gyrB*^D429N^ amplicon via electroporation as described in [37]. Transformants were selected on GCB-K plates on which 50 *μL* spots of 2- or 4 *μg*/*mL* zoliflodacin had dried. We selected transformant colonies from within the zoliflodacin zone of inhibition. The presence of the *gyrB*^D429N^ allele was confirmed by Sanger sequencing.

### Introduction of *parC*^D86N^ and *gyrB*^D429N^ into FA19

The *parC*^D86N^ mutation was inserted through a markerless transformation using the *galK*/*kanR* allelic exchange cassette as described [60]. We purchased a custom sequence from Twist Biosciences comprising the selection cassette flanked by homology surrounding the *parC* ORF. Homology arms flanking the selection cassette included the region upstream of *parC* (531 nt upstream-7 nt upstream) on one side of the cassette, and the promoter and beginning of the *parC* ORF on the other side (146 nt upstream of *parC* and the first 774 nt within *parC*). Note that the sequence 146-7 nts upstream of *parC* is repeated on each side of the cassette to ensure that the native promoters driving both *parC* and the upstream gene *gpmA* remain intact. Homology sequences were derived from the genome sequence of CCC033, an isolate that carries the *parC*^D86N^ allele [25, 32].

We amplified the selection cassette with flanking homology by PCR using primers PB_47 and PB_48. We then inserted the resulting amplicon into FA19 by electroporation as above and selected transformants on GCB-K with 70 *μ*g/mL kanamycin. We confirmed transformants by PCR and Sanger sequencing with PB_47 and PB_48. To remove the selection cassette, we performed a second-step transformation as described [60] by amplifying the homologous *parC* locus from CCC033 using the same primers and introducing it via electroporation. Markerless transformants were selected on GCB-K supplemented with 1% 2-deoxy-galactose (DOG) and confirmed by PCR and Sanger sequencing using PB_47 and PB_48.

The *gyrB*^D429N^ allele was introduced into both FA19 and FA19 *parC*^D86N^ as described above.

### Introduction of *gyrA*^S91F/A92P/D95Y^, *parE*^D437N^, and *gyrB*^D429N^ into GCGS0481 *gyrA*^91S/95D^

The *gyrA*^S91F/A92P/D95Y^ allele was amplified from HHH012 using primers AM_5 and AM_18 and introduced into GCGS0481 *gyrA*^91S/95D^ (strain number AM370), a ciprofloxacin-susceptible variant of GCGS0481 [61], via electroporation as above. Transformants were selected on GCB-K plates on which 50 *μL* spots of 2 *μ*g/mL ciprofloxacin had dried. Transformant colonies were selected from within the ciprofloxacin zone of inhibition, and the presence of the *gyrA*^S91F/A92P/D95Y^ allele was confirmed by Sanger sequencing.

The *parE*^D437N^ allele was amplified from EEE018 using primers SB_15 and SB_16 and transformed into GCGS0481 *gyrA*^91S/95D^ and GCGS0481 *gyrA*^91F/92P/95Y^ via electroporation as above. Transformants were selected on GCB-K plates on which 50 *μL* spots of 4 *μg*/*mL* gepotidacin had dried, and transformant colonies were selected from within the gepotidacin zone of inhibition. The presence of the *parE*^D437N^ allele was confirmed by Sanger sequencing.

The *gyrB*^D429N^ allele was introduced into each of these strains as described above.

### Introduction of *gyrA*^91S/92A/95D^ and *parE*^437D^ to EEE018

The wildtype *gyrA*^91S/92A/95D^ allele with an upstream kanamycin resistance marker was amplified from GCGS0481_SD (strain number YG0073), a kanamycin-resistant derivative of GCGS0481 with the *gyrA*^91S/92A/95D^ allele from FA19[37], with primers AM_5 and AM_18. We introduced the resulting amplicon into EEE018 and EEE018 *gyrB*^D429N^ by electroporation as above, and selected clones on 70 *μ*g/mL kanamycin. The presence of the *gyrA*^91S/92A/95D^ allele was confirmed by Sanger sequencing. A kanamycin-resistant clone that had retained the native *gyrA*^91F/92P/95Y^ allele due to a recombination breakpoint between the kanamycin marker and *gyrA*^91^ was retained for use as the isogenic comparator strain denoted EEE018* in table 1.

To introduce the *parE*^437D^ allele into EEE018, we constructed the pSB1 plasmid, which introduces a kanamycin resistance marker downstream of the *parE* gene. We amplified the *parE*^437D^ allele from FA19 with primers SB_25 and SB_26, the kanamycin resistance marker from pDR53[61] with primers SB_23 and SB_24, a ≈350 nt region downstream of *parE* from EEE018 with primers SB_21 and SB_22, and the plasmid backbone from pUC19 with primers SB_27 and SB_28. We assembled the PCR fragments by Gibson assembly. The kanamycin resistance marker, flanked by the end of the *parE* ORF on one side and downstream homology on the other, was amplified from pSB1 with primers SB_21 and SB_26. We introduced this amplicon into EEE018 and EEE018 *gyrB*^D429N^ via electroporation as above, and selected clones on GCB-K with 70 *μ*g/mL kanamycin. We Sanger sequenced transformants to determine whether the recombined region upstream of the kanamycin cassette included the *parE*^437D^ substitution. A kanamycin-resistant clone that retained the native *parE*^437N^ allele as a result of a recombination breakpoint between *parE*^437^ and the kanamycin cassette was retained for each transformation (EEE018 and EEE018 *gyrB*^D429N^) and was used as the isogenic comparator strain denoted EEE018** in table 1.

## Author contributions

DH, SGP, YHG conceptualized the study. DH developed and implemented the statistical methodology. SB, BB, AM, SGP designed and performed the laboratory experiments. DH, SGP, SB wrote the original draft. YHG acquired funding. DH, SGP, SB, YHG, BB, and AM discussed the results and contributed to writing, reviewing, and editing of the manuscript.

## Data, code and materials availability

The BASSGWAS package and helper scripts are available at https://github.com/dhelekal/BASSGWAS. The package versions and additional code that were used for benchmarking and experimental application are available at https://github.com/gradlab/BASSGWAS-Experiments. Benchmark datasets and annotated GWAS outputs were deposited at https://doi.org/110.5281/zenodo.22049989.

## Supplementary Materials

### Supplementary Text

#### Sparse Whole-Genome Regression Models

We based our model on the seminal BSLMM introduced by [26]. As in the BSLMM, we represented phenotypic variation as the sum of two components [26]. The first component accounts for the impact of variants with large effect sizes. We used *β* to denote the vector of coefficients corresponding to the effects of such variants. The model assumes that only a small fraction of all variants have large effects. To impose this assumption, the model uses a spike-and-slab prior [49, 63] that constrains only a fraction of elements in *β* to be non-zero. The second component accounts for the total effect of the genetic background. We used *u* to denote the vector coefficients corresponding to this effect. Given a matrix *X* that encodes the presence and absence of genetic variants and a matrix *K* that encodes the population structure, the linear predictor *ψ* for strain *j, ψ*_*j*_, is given by:

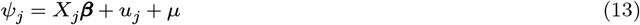

Here, *μ* denotes the intercept. Additionally, we introduced two parameters, 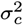 and 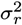, that govern the effect scales for variants with large effects and for the genetic background, respectively:

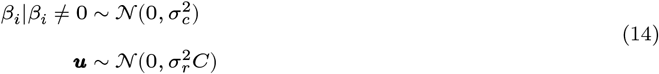

The linear predictor is connected to the observations using a binary Probit likelihood [27]

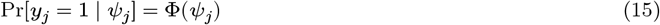

Here, Φ denotes the cumulative distribution function of the standard normal distribution. The effect scale of the genetic background 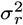 is estimated as part of the model. The effect scale for variants with large effects 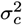 can be estimated as a free parameter or treated as a fixed constant; we implemented our model both ways and found close agreement between these approaches for a range of benchmarking tests (see e.g. Figures 3 vs S2).

#### Benchmark Phenotype Description

High-level tetracycline resistance (MIC ≥ 8 *μ*g/mL) is conferred by the carriage of the plasmid-borne gene *tetM* [32] with high phenotypic penetrance. Ciprofloxacin resistance (MIC ≥ 1 *μ*g/mL) is a polygenic trait that requires point mutations in both *gyrA* and *parC* [32] to attain the resistant phenotype. Azithromycin non-susceptibility (MIC ≥ 2 *μ*g/mL) can be caused by genetically diverse recombinant mosaic alleles and insertions/deletions in the *mtrCDE* operon as well as mutations in 23S ribosomal rRNA [31].

#### Regression Model Validation

For tetracycline resistance, *tetM* was identified with high probability (PIP = 1) by all three types of posterior summary analysis: individual unitig analysis identified a single variant indicating the presence of *tetM* (Tables S3); analysis of genomic regions identified a locus on the conjugative plasmid surrounding the *tetM* insertion site that is known to be unique to the plasmids that carry *tetM* [64] (Figure 2); and genomic annotation analysis identified *tetM* and several surrounding adjacent genes that are linked to the presence of *tetM* (one gene upstream and four genes downstream) (Figure 2). For ciprofloxacin resistance, the known resistance loci in *gyrA* and *parC* were identified by all three methods with high probability (Figure 2) (Tables S3). The modestly lower PIP for *parC* variants from the individual unitig analysis (PIP=0.82) reflects decreased sensitivity of this approach for more variable loci, as multiple alternative *parC* genotypes contribute to the ciprofloxacin resistance phenotype [65]. This principle was demonstrated most clearly for azithromycin non-susceptibility: while the involvement of 23S rRNA variants and variants in the *mtrCDE* operon were identified by genomic annotation analysis with high confidence, the genomic region analysis identified the 23S rRNA region with high confidence but the region corresponding to the *mtrCDE* operon only with PIP less than 0.4 (Figure 2). This apparent discrepancy results from unitigs corresponding to divergent mosaic *mtr* alleles failing to map to a single reference genome. Similarly, individual unitig analysis successfully identified a single dominating variant at the 23S rRNA locus (Tables S3), but no individual variant mapping to the *mtrCDE* operon exceeded PIP of 0.5 (Tables S3), again due to high diversity in this locus. These results demonstrate that multiple posterior summaries can contribute to variant identification.

#### BASS-GWAS for the genetic basis of zoliflodacin/gepotidacin cross-resistance

7 batches of 4 strains each were tested for zoliflodacin/gepotidacin cross resistance, defined as a gepotidacin MIC increase of >4-fold after introduction of the zoliflodacin resistance mutation *gyrB*^D429N^. We selected these strains from our laboratory’s existing collection of *N. gonorrhoeae*, which includes strains from a variety of published sources spanning different time periods and countries [32, 66–69]. A list of tested strains and their phenotypic results is presented in (table S1). One isolate (Ng_188) was selected in two rounds but failed to provide phenotypic information both times. When it was first selected, we were unable to generate a viable *gyrB*^D429N^ clone and proceeded to strain selection for the next batch. When it was selected again, we failed to introduce the *gyrB*^D429N^ allele after three attempts. Therefore, we censored this strain from selection in subsequent batches.

##### 1 − 1/*e* Optimality of EIG Optimization

###### Definition 1

*A function f* : *P*(Ω) ↦ ℝ *is called monotone if for all A* ⊆ *B* ⊆ Ω: *f*(*A*) ≤ *f*(*B*)

For sets *A, B* with *A* ∩ *B* = ∅ we have

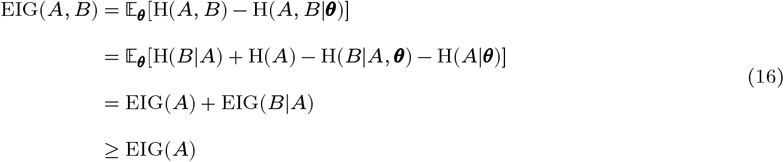

Since EIG(·)≥ = 0. Hence, the EIG is monotone.

###### Definition 2

*A function f* : *P*(Ω) ↦ ℝ *is called submodular if for all S* ⊂ Ω *and distinct Y*_1_, *Y*_2_ ∉ *S: f*(*S* ∪ *Y*_1_ ∪ *Y*_2_) + *f*(*S*) ≤ *f*(*S* ∪ *Y*_1_) + *f*(*S* ∪ *Y*_2_)

We begin by expanding the definition of EIG:

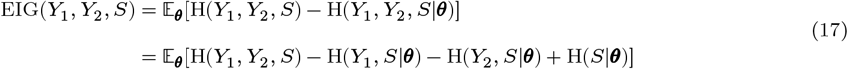

The second equality follows from the conditional independence of *X, Y, S* given *θ*. Further, expanding, we conclude that:

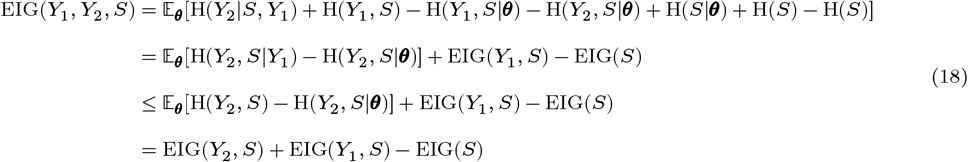

This matches the definition of a submodular function. Therefore, the EIG is a monotone, submodular function. From [56] we conclude that the greedy maximization algorithm is 1 − 1/*e* optimal.

#### Impact of Batch Size

Let *S* be a set of experiments that maximizes the current incremental EIG. Partition *S* into two non-overlapping subsets of experiments *A* ∪ *B* = *S, A* ∩ *B* = ∅. Suppose that we have observed the realization of experiments that make up *A*, i.e., *A* = *a*. Using the same decomposition as in (equation 16), under the assumption that the underlying model is correct, we conclude that incorporating the information *A* = *a* and replacing *B* with a new design *B*^∗^ can only increase the EIG:

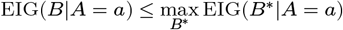

Therefore, we expect that the per-sample efficiency can only decrease as the batch size increases.

#### MCMC Sampling

Regression models involving the spike-and-slab prior are notoriously challenging to sample from. To make this problem tractable, we used weighted Tempered Gibbs Sampling (wTGS) of [52], and its extension to data augmentation schemes [53]. For discrete observations, wTGS is not directly applicable because it requires a conjugate likelihood to compute importance weights. WTGS is effectively an informed random-scan sampling scheme: First, a parameter index is chosen from an appropriate distribution; second, the corresponding parameter is updated using a tempered conditional. Thus, wTGS can be extended to data augmentation schemes and non-conjugate hyperparameters by adding an index for all such parameters and updating them by sampling from an untempered conditional [53]. We adopt this strategy using data augmentation for probit models [27].

Throughout this section, we abuse notation and suppress the posterior distribution’s dependence on the strain index variables *ξ*_1:*N*_. First, we obtain the parameter marginals needed to implement the wTGS-based sampling scheme. *π*(*y*_1:*N*_ |*z*) can be expressed as:

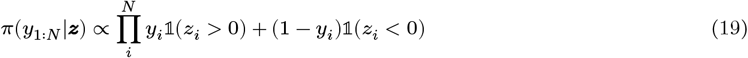

Next coefficients *β, u, μ* can be marginalized out using standard properties of multivariate normal distributions to obtain the likelihood *π*(*z*|*γ, σ*_*c*_, *σ*_*r*_) = ∫ *π*(*z*|*β, u, μ*)*π*(*β*|*γ, σ*_*c*_)*π*(*u*|*σ*_*r*_)*π*(*μ*) d*β*d*u*d*μ*:

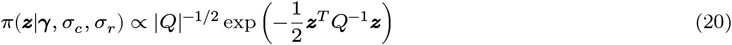

Where 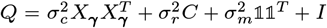, and *X* refers to *X* restricted to the column indices for which *γ*_*j*_ = 1. Note that *Q*^−1^ can be computed efficiently using the Woodbury identity.

*π*(*γ*) can be expressed as:

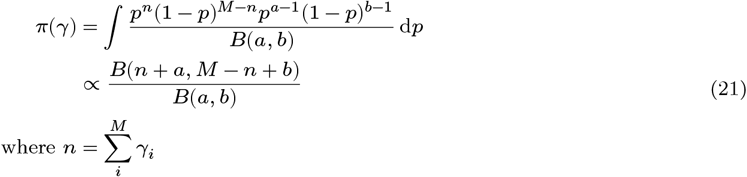

To improve mixing, we adopted a marginal data augmentation scheme based on [54], whereby an unidentifiable scale parameter is introduced, together with a shift move that shifts the random utilities *z* using an auxiliary parameter. We first introduced a scale parameter *ζ* and defined the scaled utilities 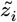 as:

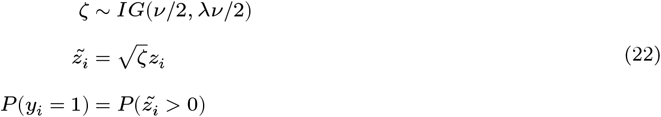

Next, based on [53] we partition the parameters of the augmented posterior as 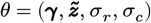. The wTGS scheme for this distribution then admits the following mixture representation

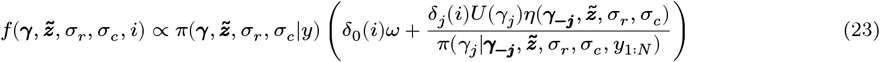

Here, *U*(·) denotes the uniform distribution on {0, 1} and *ω* is a hyperparameter that governs the frequency of updates for (*z, σ*_*c*_, *σ*_*r*_). Before sampling, we run a single short chain to identify *ω* so that the frequency of updates for (*z, σ*_*c*_, *σ*_*r*_) approximates a target frequency. We set the target frequency to 0.25. We set the weight 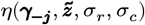 as in [52]:

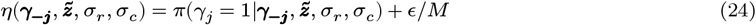

We used *ϵ* = 5. To obtain 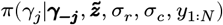, we note that:

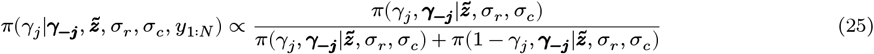

Here, 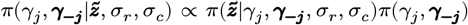. Moreover, the marginal distribution 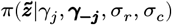 can be obtained in a closed form, using the conjugacy between the multivariate normal and the inverse-gamma distribution:

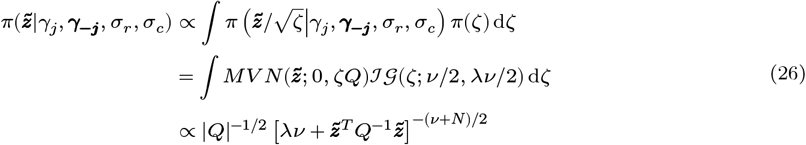

WTGS proceeds by sampling *i* proportional to its marginal probability followed by an update using a markov kernel that is invariant with respect to the tempered *i*−th conditional. For *i* ≠ 0, we set the tempered conditional to the uniform distribution on {0, 1} and we use a metropolized kernel that has an acceptance probability of 1:

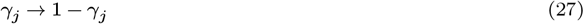

See [52, 53] for details of the wTGS sampler and its extension to data-augmentation schemes.

The move corresponding to *i* = 0 varied between the version with varying large effect scale and the version with fixed effect scale.

##### Varying large effect scale

We use a sequence of moves to update each of 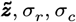:

1. We sample 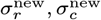 using an RWM kernel with a reflecting boundary:

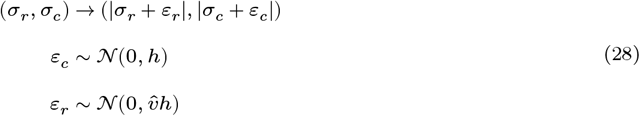

Here 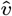 is an estimate of the variance ratio var(*r*)/var(*c*) obtained during burn-in, and *h* is a step-size optimized during burn-in. The corresponding acceptance probability for this move is:

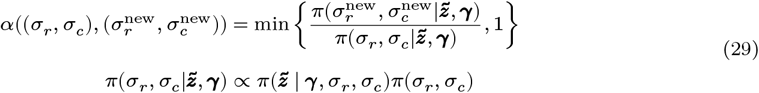

We found that the posterior distribution of *σ*_*r*_, *σ*_*c*_ can be heavy tailed, impeding sampling. We therefore iterated this move multiple times with a monotonically increasing step-size *h*, performing an accept/reject step after every iteration.
2. We sample the augmented scale parameter *ζ* from the respective conjugate conditional 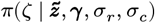 [54]:

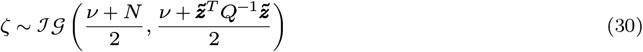

and set 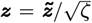. Next, we use the Gibbs data augmentation kernel for the Probit model [27] to sample *z*^new^:
  a. We draw *β, u, μ* from its multivariate normal conditional *π*(*β, u, μ*|*z, γ, σ*_*r*_, *σ*_*c*_) and use *β, u, μ* to compute the linear predictor *ψ* (Equation 1).
  b. We sample 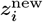 from a truncated normal distribution 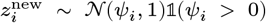 if *y*_*i*_ = 1 or 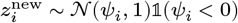 if *y*_*i*_ = 0.
3. We use the shift move of [54] to shift *z*^new^
4. We sample a new scale parameter 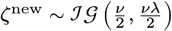 and set 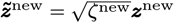

#### Fixed large effect scale

We use a sequence of moves to update each of 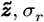:

1. We sample the augmented scale parameter *ζ* from the respective conjugate conditional 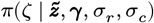 [54]:

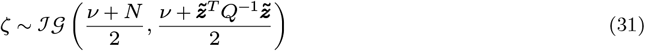

and set 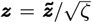. Next, we use the Gibbs data augmentation kernel for the Probit model [27] to sample *z*^new^:
  a. We draw *β, u, μ* from its multivariate normal conditional *π*(*β, u, μ*|*z, γ, σ*_*r*_, *σ*_*c*_) and use *β, u, μ* to compute the linear predictor *ψ* (Equation 1).
  b. We sample 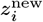 from a truncated normal distribution 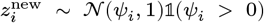 if *y*_*i*_ = 1 or 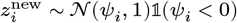 if *y*_*i*_ = 0.
2. We sample 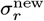 using an RWM kernel with a reflecting boundary:

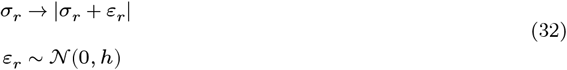

Here *h* is a step-size optimized during burn-in. The corresponding acceptance probability for this move is:

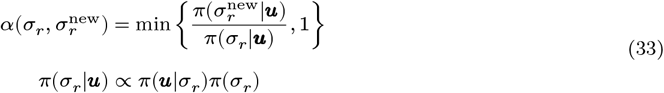

We iterate this move multiple times with a monotonically increasing step-size *h*, performing an accept/reject step after every iteration.
3. We use the shift move of [54] to shift *z*^new^
4. We sample a new scale parameter 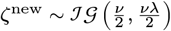 and set 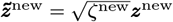

### Supplementary Figures

**Figure S1:**
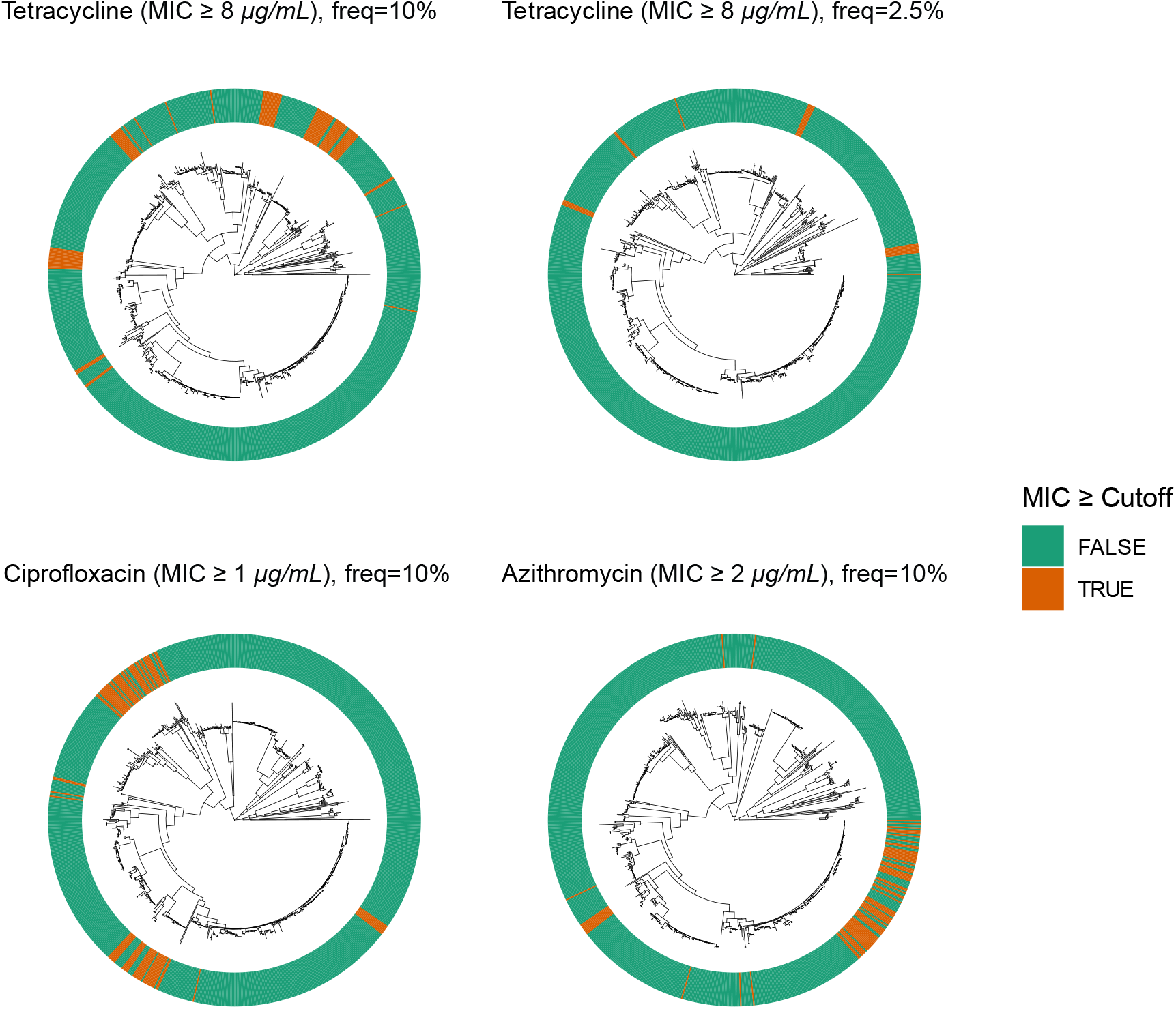
Phylogenetic Distribution of Phenotypes in Benchmark Datasets. The color strip depicts the distribution of the phenotypic resistance across the phylogeny.

**Figure S2:**
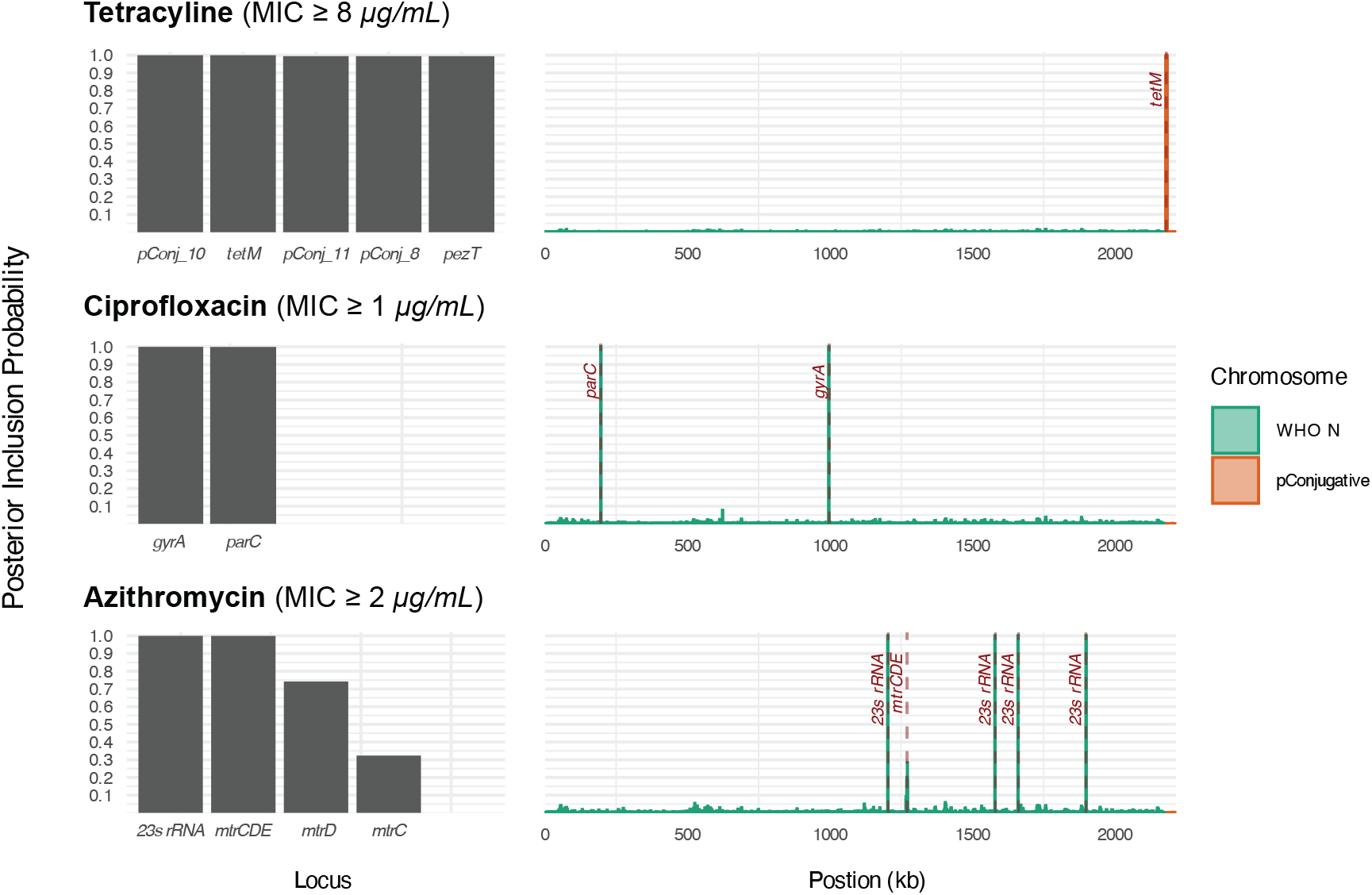
Validation of the Whole Genome Regression Model With Fixed Large Effect Scale Using Three Phenotypes. Top: Tetracycline resistance. Middle: Ciprofloxacin resistance. Bottom: Azithromycin non-susceptibility. Left column: posterior inclusion probabilities (PIPs) for annotated genomic elements; shown: all annotated loci with PIP exceeding 0.25. Right column: PIPs for genomic regions, calculated for 1 kb overlapping windows spaced 500 bp apart. Locations of known causal loci are labeled using dashed lines and text. Shown: The WHO N chromosome and the conjugative plasmid. The gene *pezT* and the hypothetical proteins *pConj_8,10,11* flank the *tetM* insertion site and have perfect linkage with *tetM. mtrCDE* is the MtrCDE efflux pump operon.

**Figure S3:**
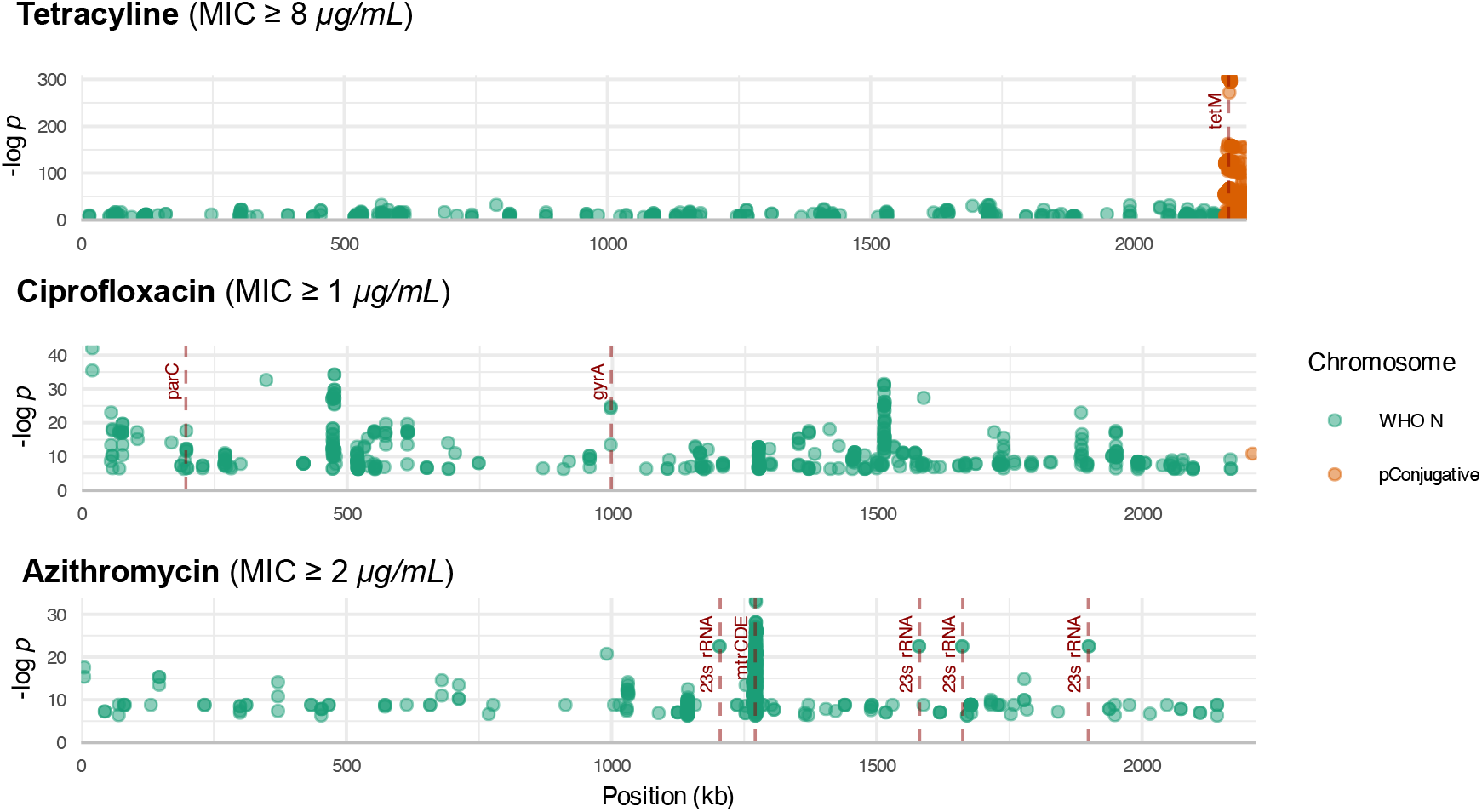
Mapping of Associated Variants Using a Marginal LMM. Points represent significantly associated *unitigs* mapped to the *WHO N* reference. Significance testing was performed using *pyseer*. The significance level was set to *α* = 0.95 after a multiple-testing correction. Annotations in red indicate the locations of known causal loci.

**Figure S4:**
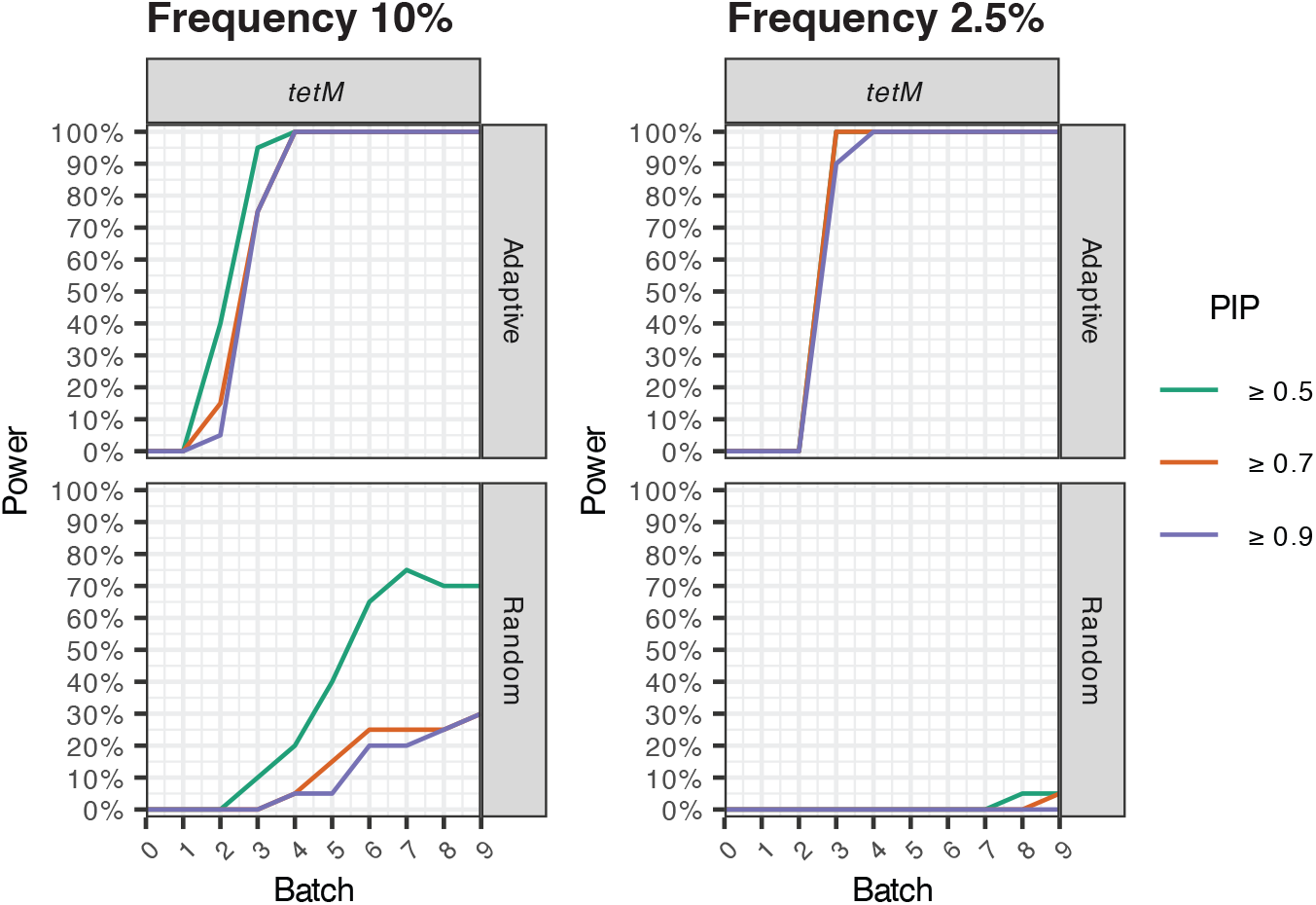
Tetracycline Resistance Power Comparison. The figure depicts the percentage of replicates (20 simulated experiments) for which the PIP of the causal gene (*tetM*) exceeded a given threshold after assaying a given number of batches. Batch 0 consists of the initial randomly selected samples. Batch size of 8. The model with a varying large effect scale was used.

**Figure S5:**
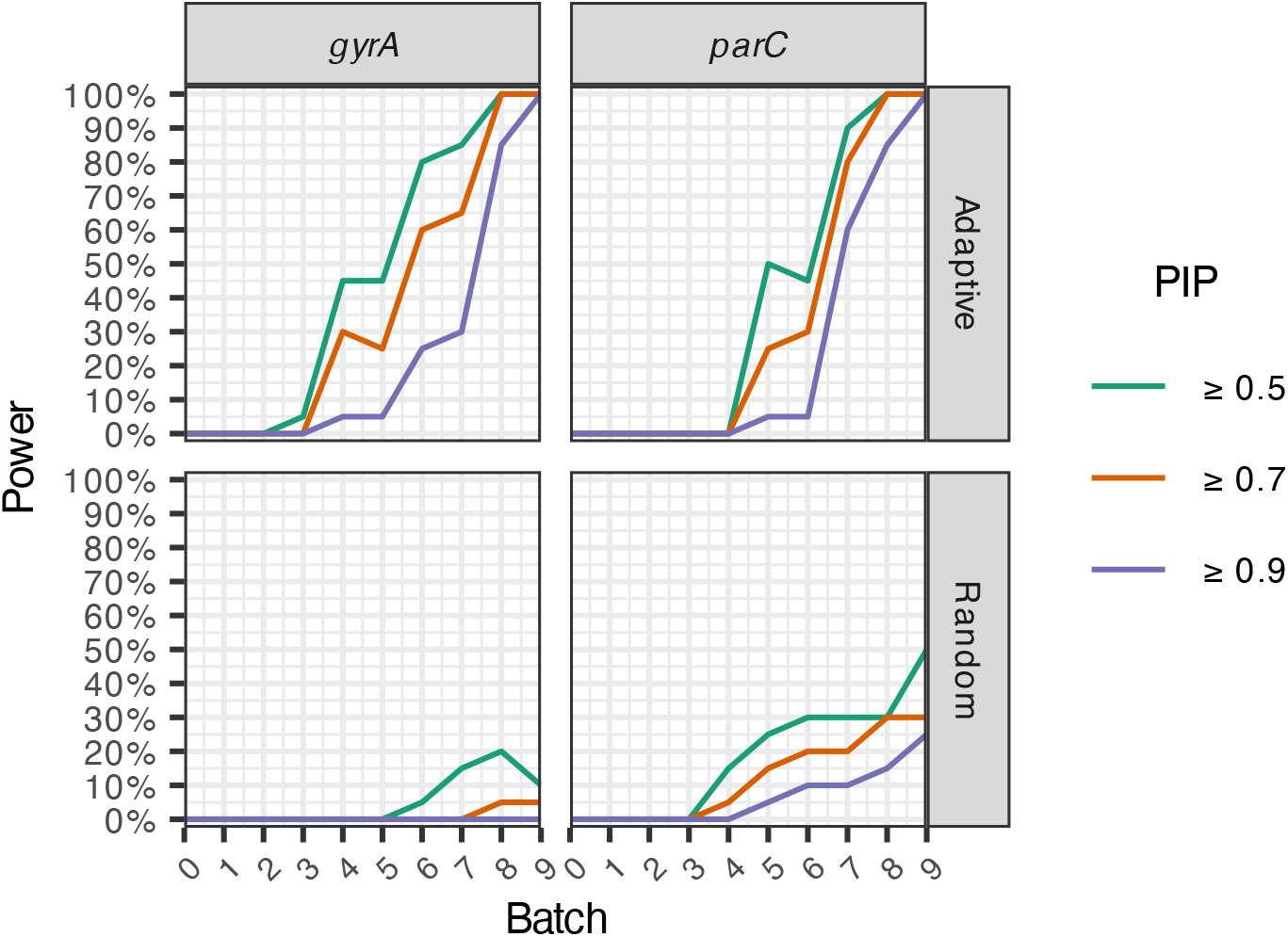
Ciprofloxacin Resistance Power Comparison. The figure depicts the percentage of replicates (20 simulated experiments) for which the PIP of the causal genes *gyrA* and *parC* exceeded a given threshold after assaying a given number of batches. Batch 0 consists of the initial randomly selected samples. Batch size of 8. The model with a varying large effect scale was used.

**Figure S6:**
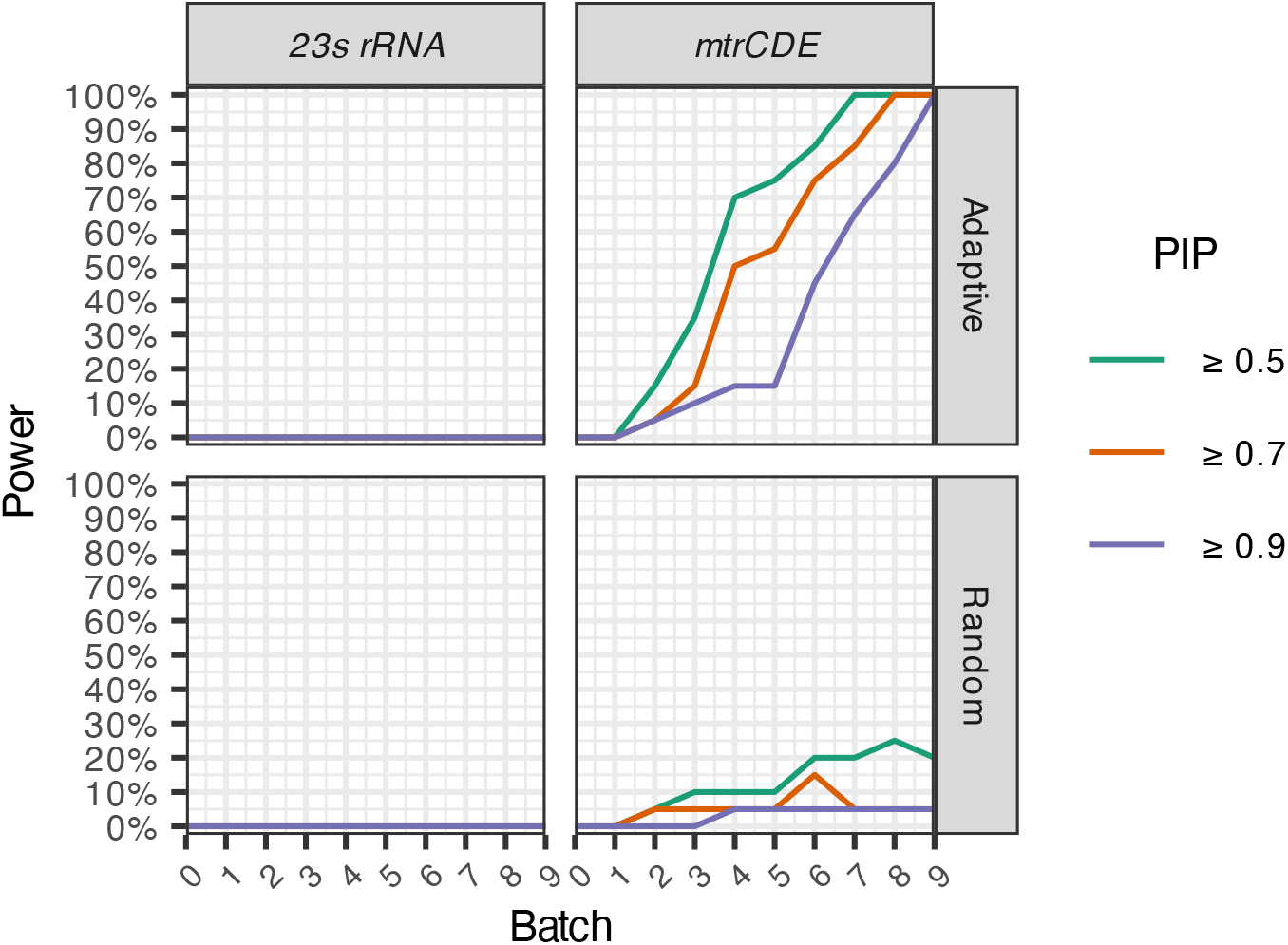
Azithromycin Resistance Power Comparison. The figure depicts the percentage of replicates (20 simulated experiments) for which the PIP of the causal genes *23s rRNA* and *mtrCDE* exceeded a given threshold after assaying a given number of batches. Batch 0 consists of the initial randomly selected samples. Batch size of 8. The model with a varying large effect scale was used.

**Figure S7:**
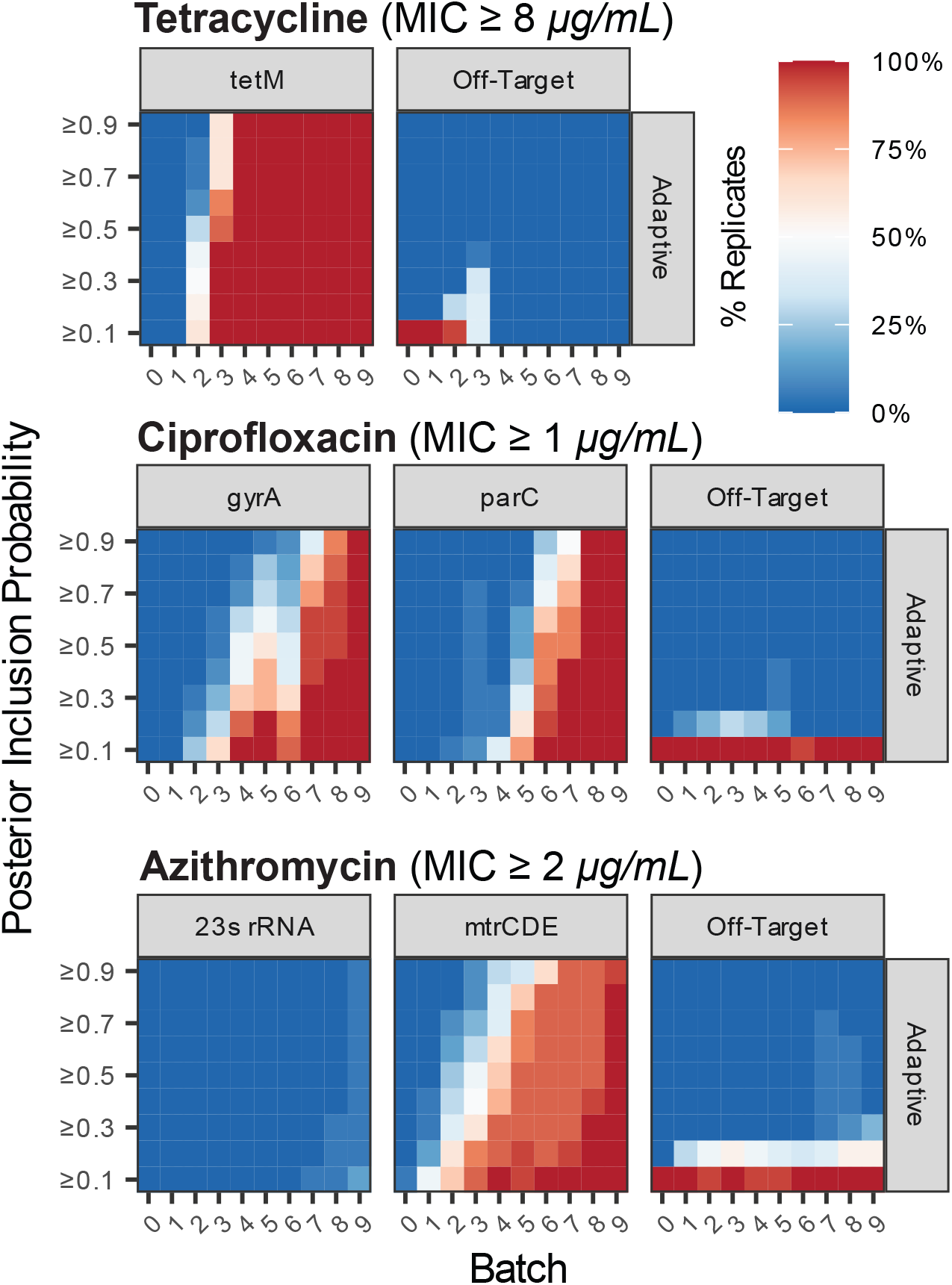
Performance of Adaptive Sampling for the Model With Fixed Large Effect Scale. The heatmap depicts the percentage of replicates (20 simulated experiments) for which the PIP of an annotated gene exceeded each threshold after assaying a given number of batches (8 strains per batch), highlighting the PIPs of known causal loci and the highest PIP among all remaining annotations (denoted Off-Target). Causal locus for tetracycline resistance: *tetM*. Causal loci for ciprofloxacin resistance: *gyrA* and *parC*. Causal loci for azithromycin non-susceptibility: the *mtrCDE* operon and 23S rRNA. Batch 0 consists of the initial randomly selected samples.

**Figure S8:**
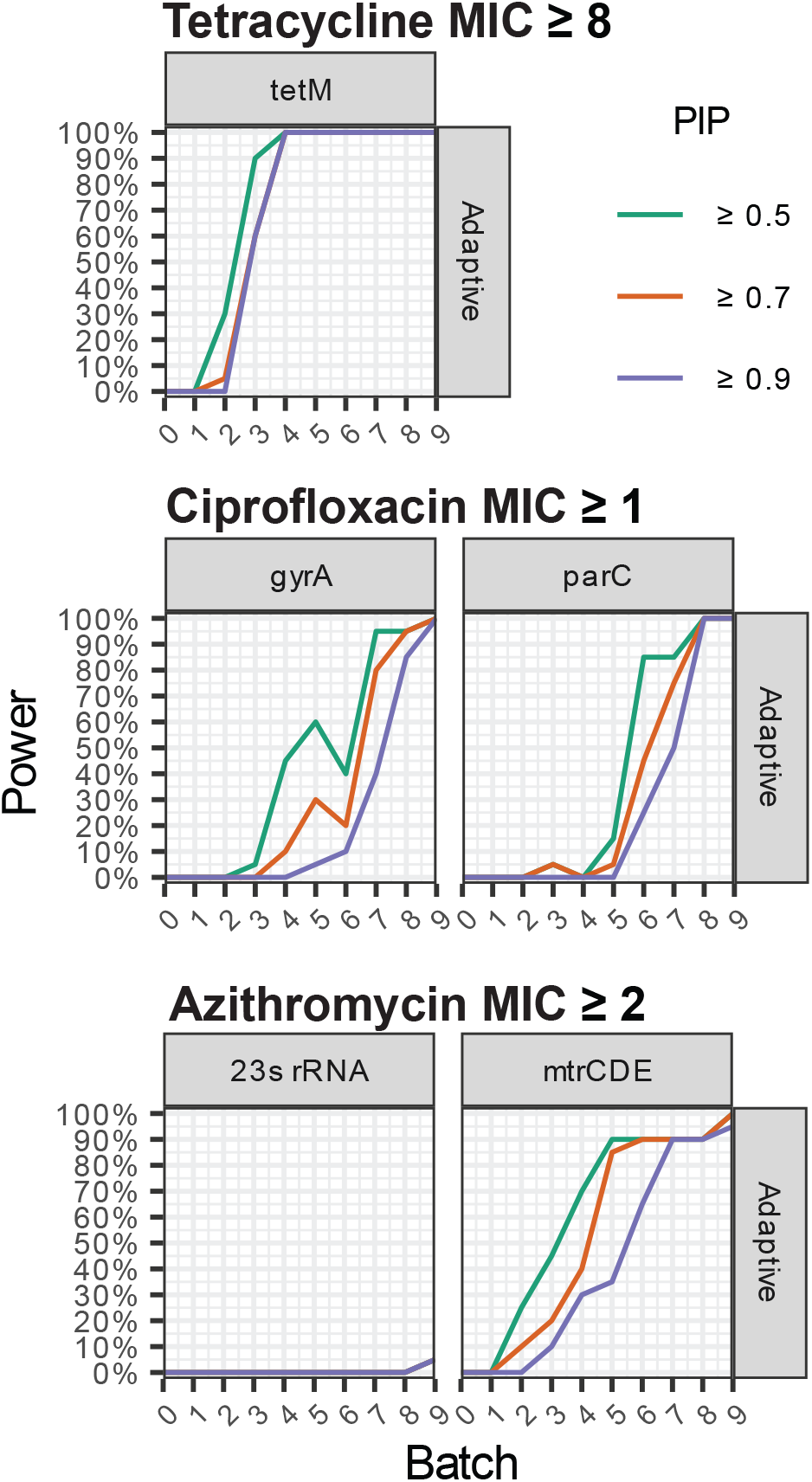
Power Analysis of Adaptive Sampling for the Model With Fixed Large Effect Scale. The heatmap depicts the percentage of replicates (20 simulated experiments) for which the PIP of an annotated gene exceeded a given threshold after assaying a given number of batches (8 strains per batch), highlighting the PIPs of known causal loci and the highest PIP among all remaining annotations (denoted Off-Target). Causal locus for tetracycline resistance: *tetM*. Causal loci for ciprofloxacin resistance: *gyrA* and *parC*. Causal loci for azithromycin non-susceptibility: the *mtrCDE* operon and 23S rRNA. Batch 0 consists of the initial randomly selected samples.

**Figure S9:**
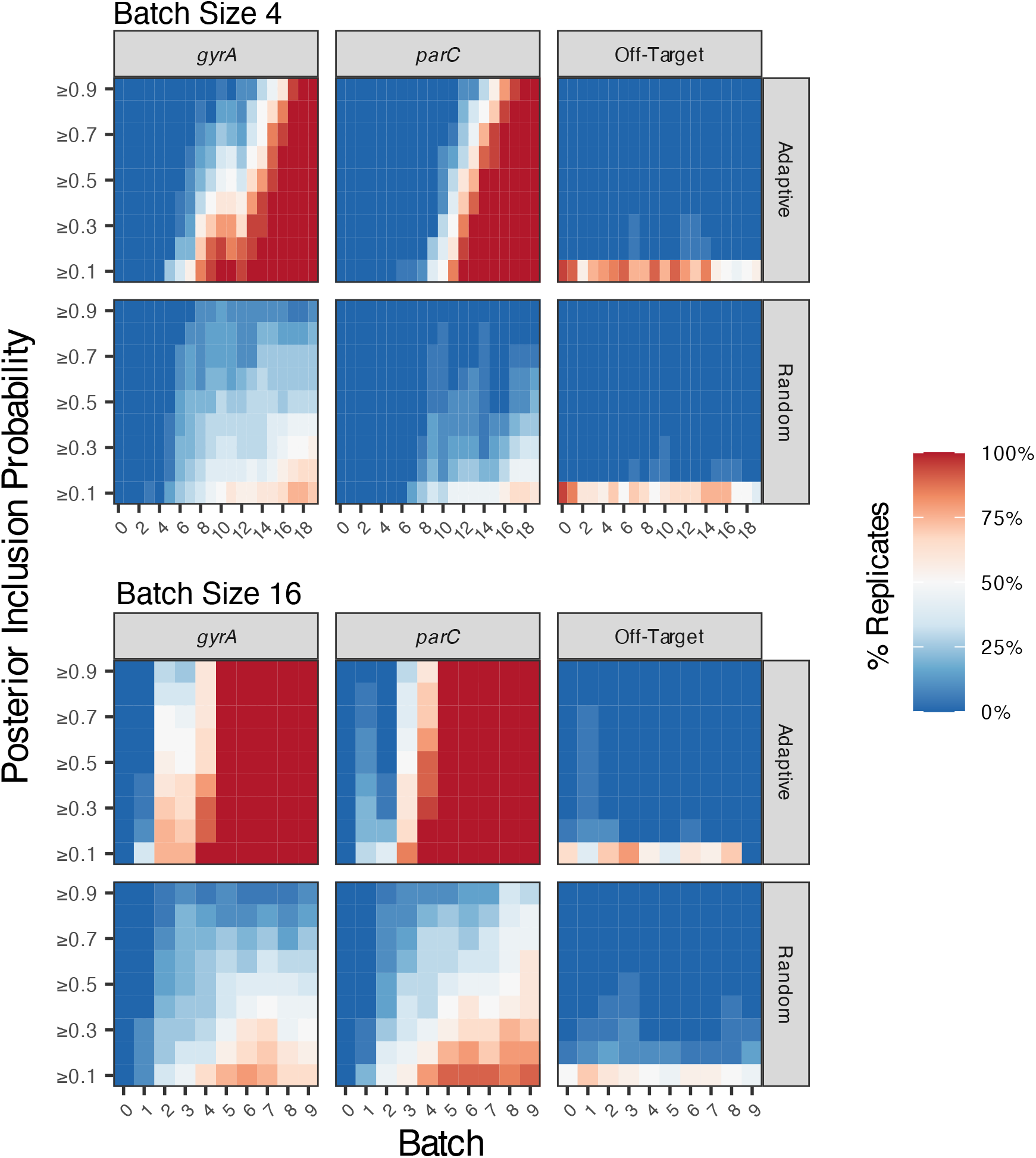
A Comparison of Adaptive and Random Sampling With Batch Size of 4 and 16 Using Ciprofloxacin Resistance Dataset. The heatmap depicts the percentage of replicates (20 simulated experiments) for which the PIP of an annotated gene exceeded each threshold after assaying a given number of batches. The figure displays the PIPs of annotations of known causal loci for ciprofloxacin resistance (*gyrA* and *parC*) and the highest PIP among all remaining annotations (denoted as *Off-Target*). Batch 0 consists of initial randomly selected samples equal to the batch size. Top: Batch size of 4. Bottom: Batch size of 16. The model with a varying large effect scale was used.

**Figure S10:**
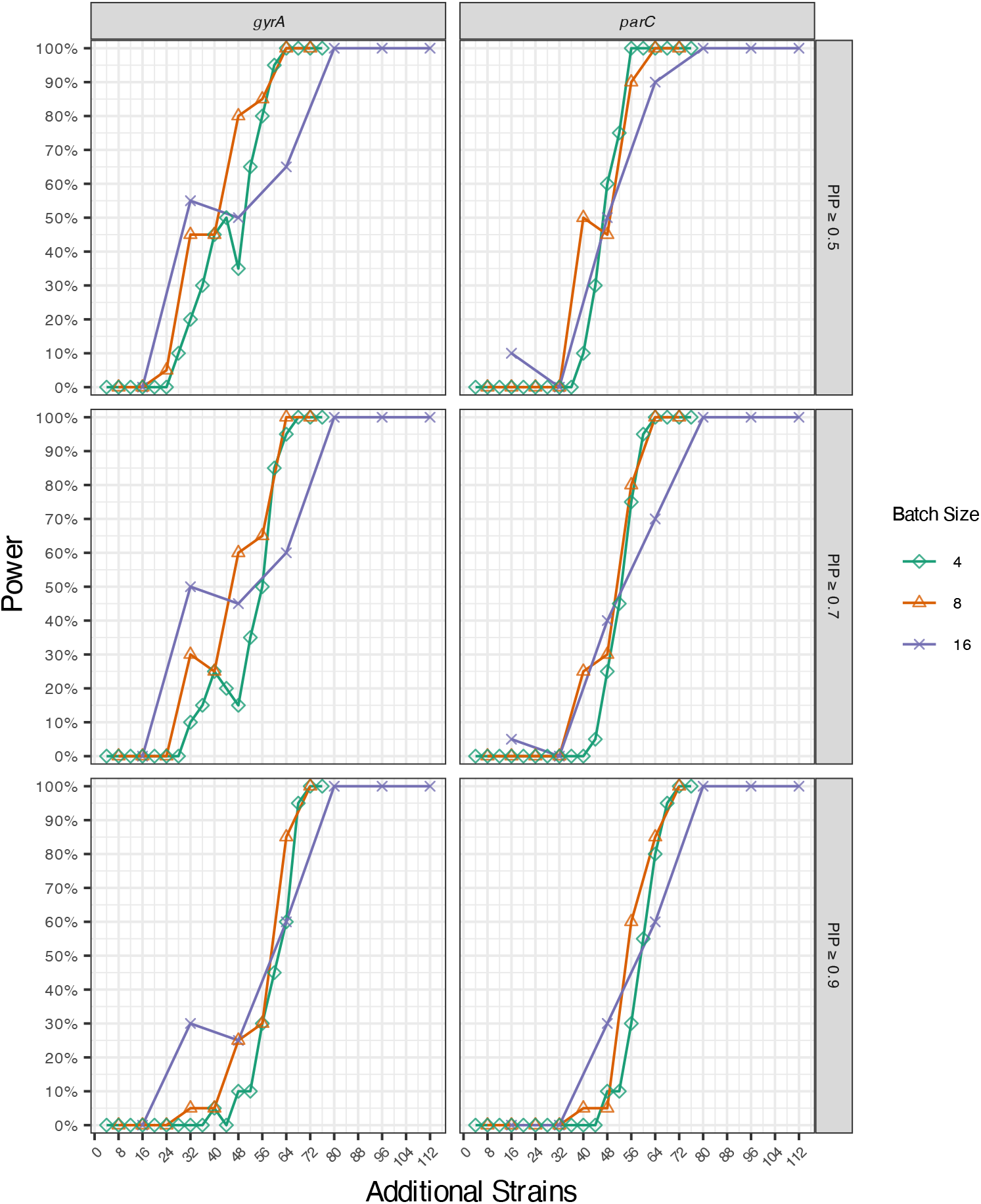
The Impact of Batch Size on the Per-Sample Efficiency of Adaptive Sampling for Ciprofloxacin Resistance. The figure depicts the percentage of replicates (20 simulated experiments) for which the PIP of an annotated causal gene exceeded 0.5 (top), 0.7 (middle), or 0.9 (bottom) after assaying a given number of additional strains. Assays were performed in batches of 4, 8, or 16 strains at a time. The model with a varying large effect scale was used.

**Figure S11:**
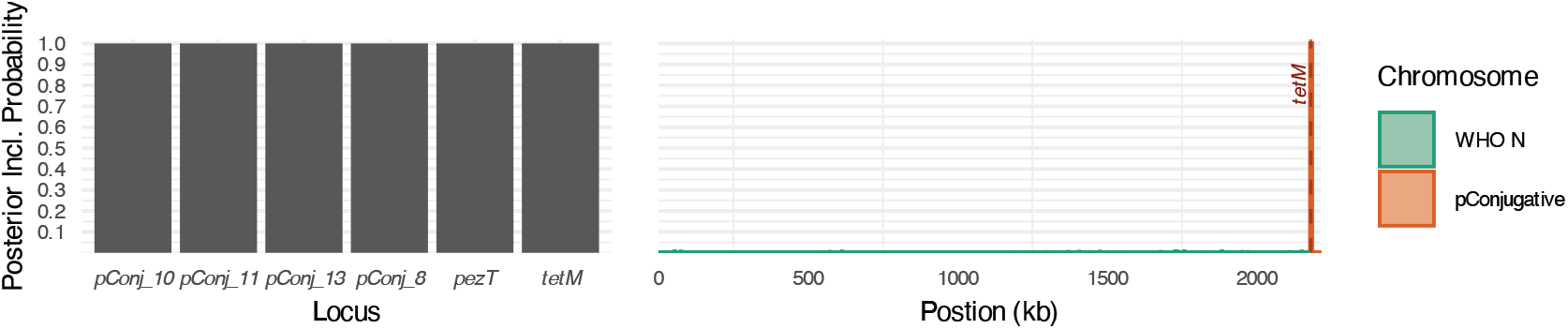
Validation of The Whole Genome Regression Tetracyline Resistance Dataset with 2.5% Prevalence. Left column: posterior inclusion probabilities (PIPs) for annotated genomic elements with PIP exceeding 0.25. Right column: PIPs for genomic regions. PIPs are calculated for 1 kb overlapping windows, spaced 500 bp apart. Locations of known causal loci are labeled in red. The model with a varying large effect scale was used. Shown: The WHO N chromosome and the conjugative plasmid.

**Figure S12:**
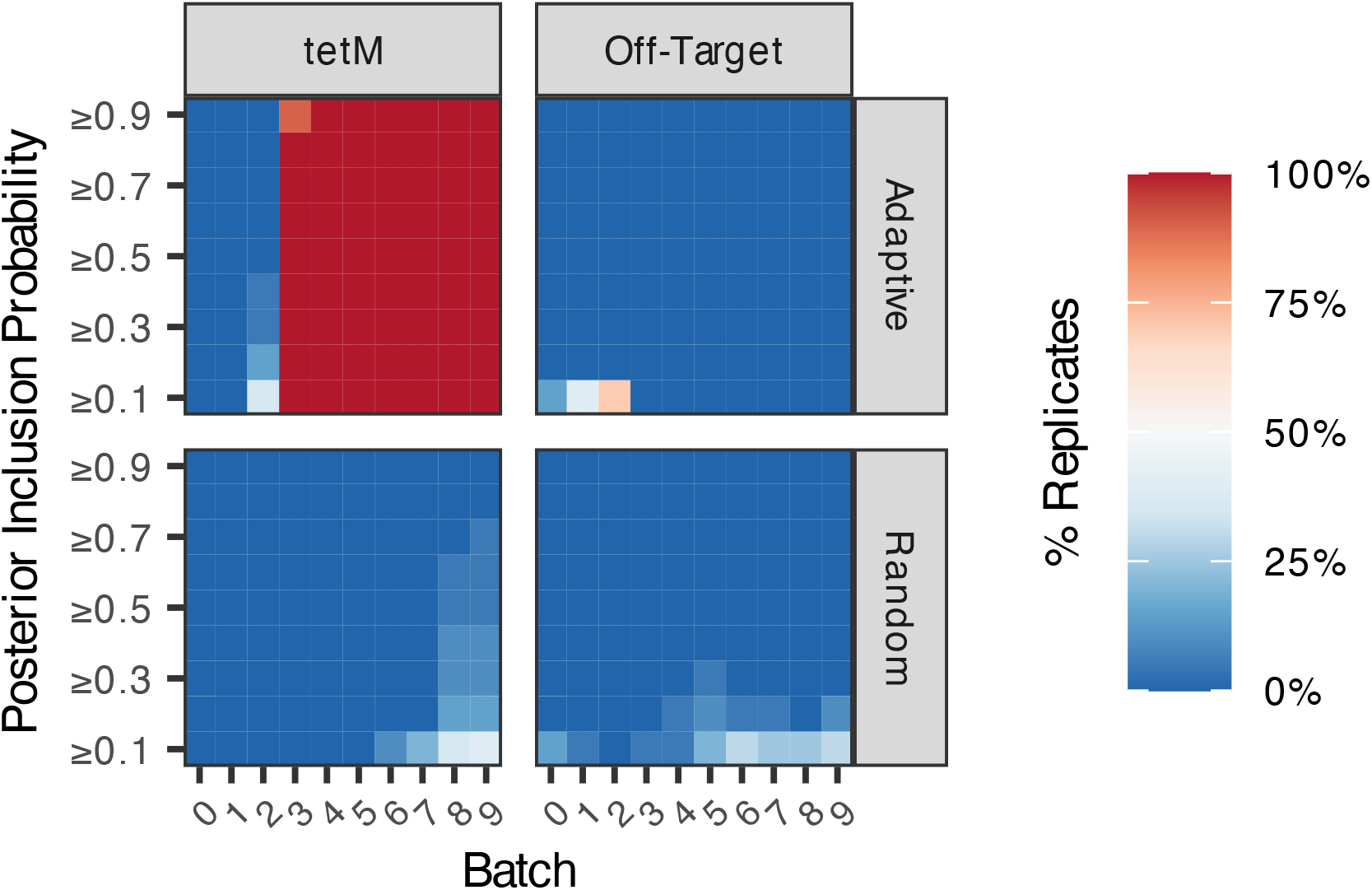
A Comparison of Adaptive and Random Sampling Using Tetracyline Resistance Dataset with 2.5% Prevalence. The heatmap depicts the percentage of replicates (20 simulated experiments) for which the PIP of an annotated gene exceeded each threshold after assaying a given number of batches, here showing the PIPs of annotations of known causal loci (*tetM*) and the highest PIP among all remaining annotations (denoted as *Off-Target*). Batch 0 consists of the initial randomly selected samples. Batch size of 8. The model with a varying large effect scale was used.

**Figure S13:**
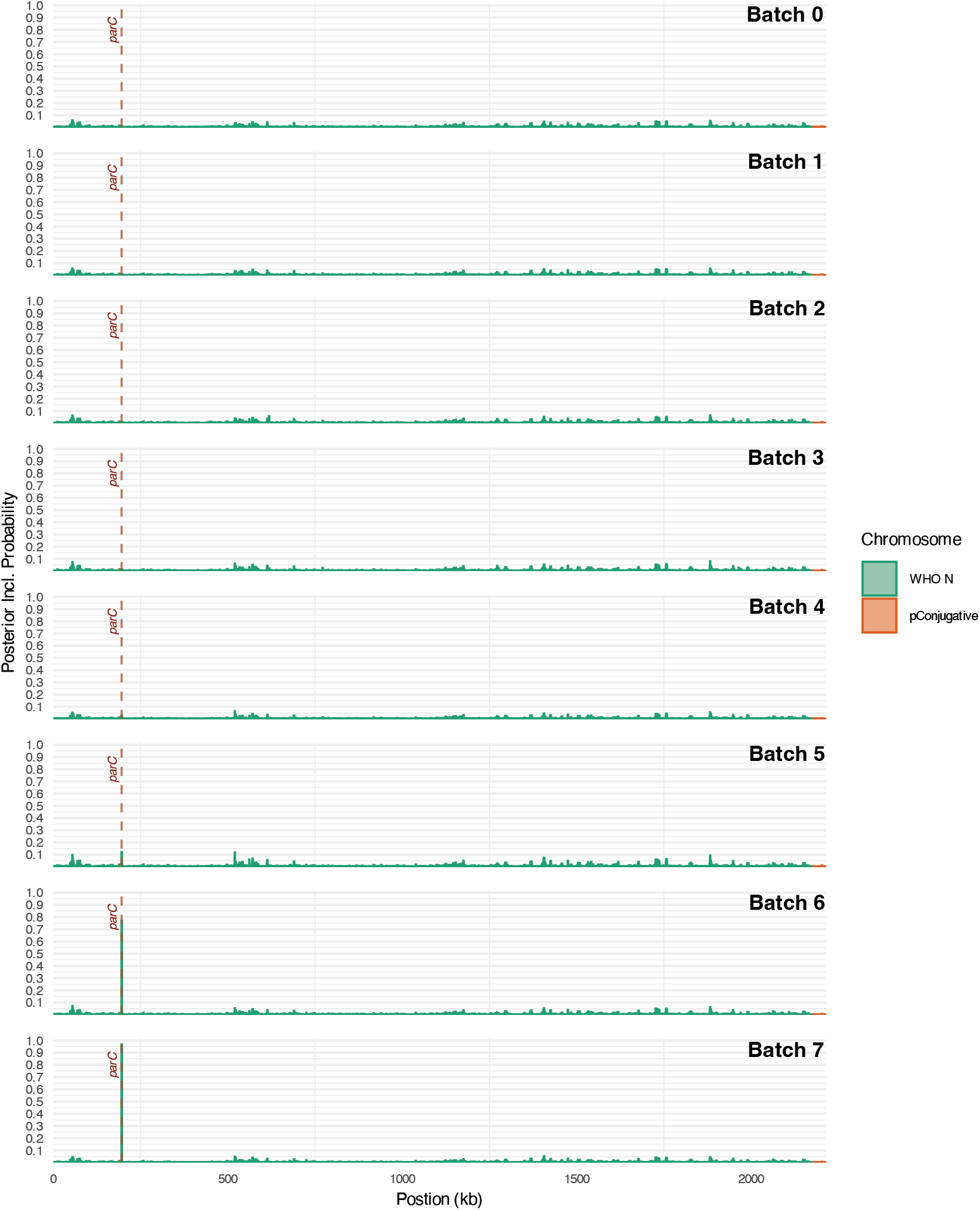
Posterior Inclusion Probabilities of Genomic Regions Across Multiple Experiments. PIPs are calculated for 1 kb overlapping windows, spaced 500 bp apart. The position of *parC* is annotated in red. Shown: The WHO N chromosome and the conjugative plasmid.

**Figure S14:**
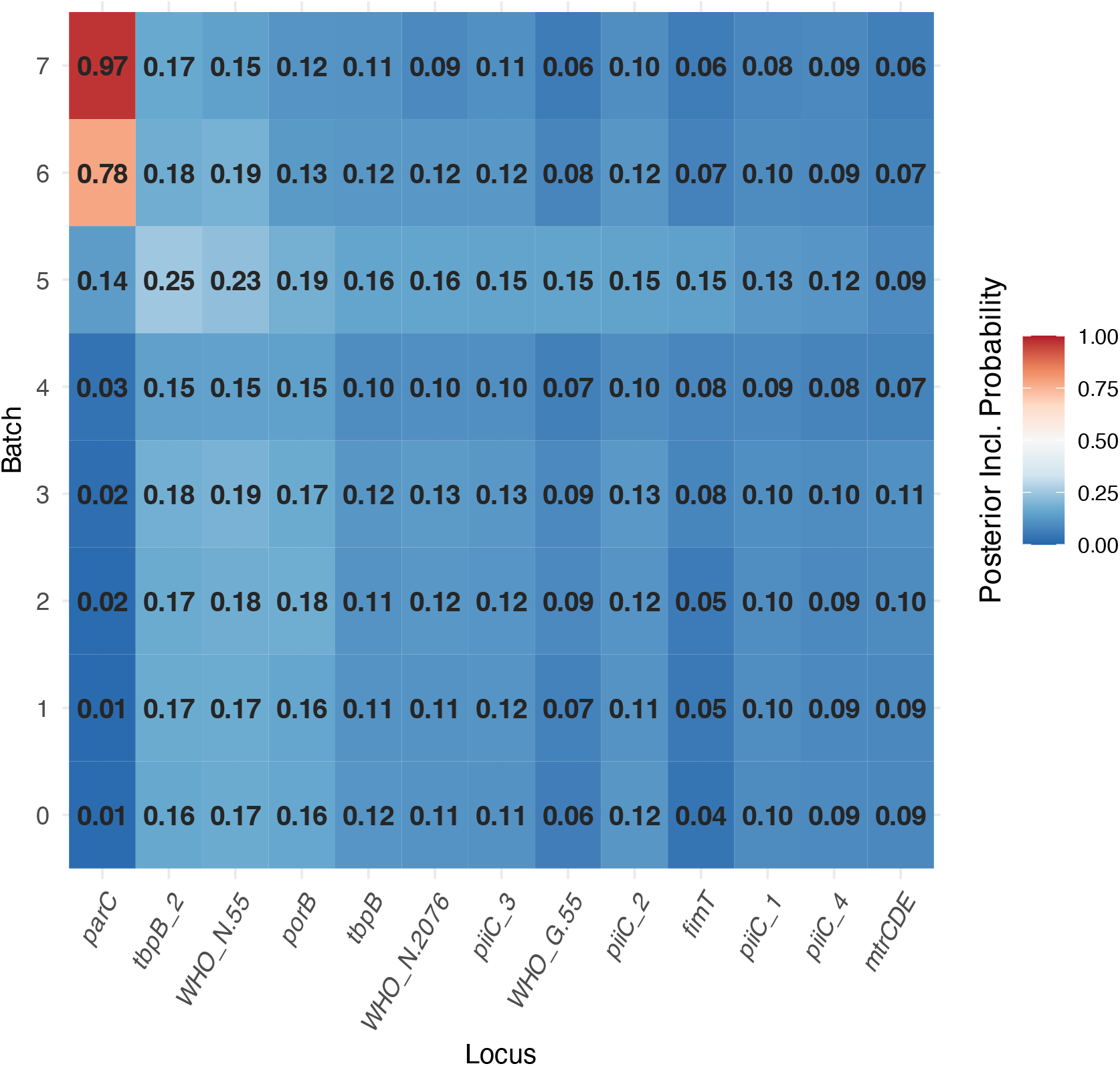
Posterior Inclusion Probabilities of Genomic Annotations Across Multiple Experiments. A heatmap depicting the evolution of PIPs for select annotations over the course of successive experiments. Columns correspond to annotations for which the corresponding PIP exceeded 0.1 at any point. Rows correspond to experimental batches. PIPs are indicated in bold.

### Supplementary Tables

Table S1: **Identifiers, MICs, and References for Strains Generated or Phenotyped as a Part of This Study**.

Table S2: **Accessions and Phenotypes of Strains Included in Benchmark Datasets**.

Table S3: **PIPs of Individual Variants for Whole-Genome Regression Model Validation**.

**Table S4:**
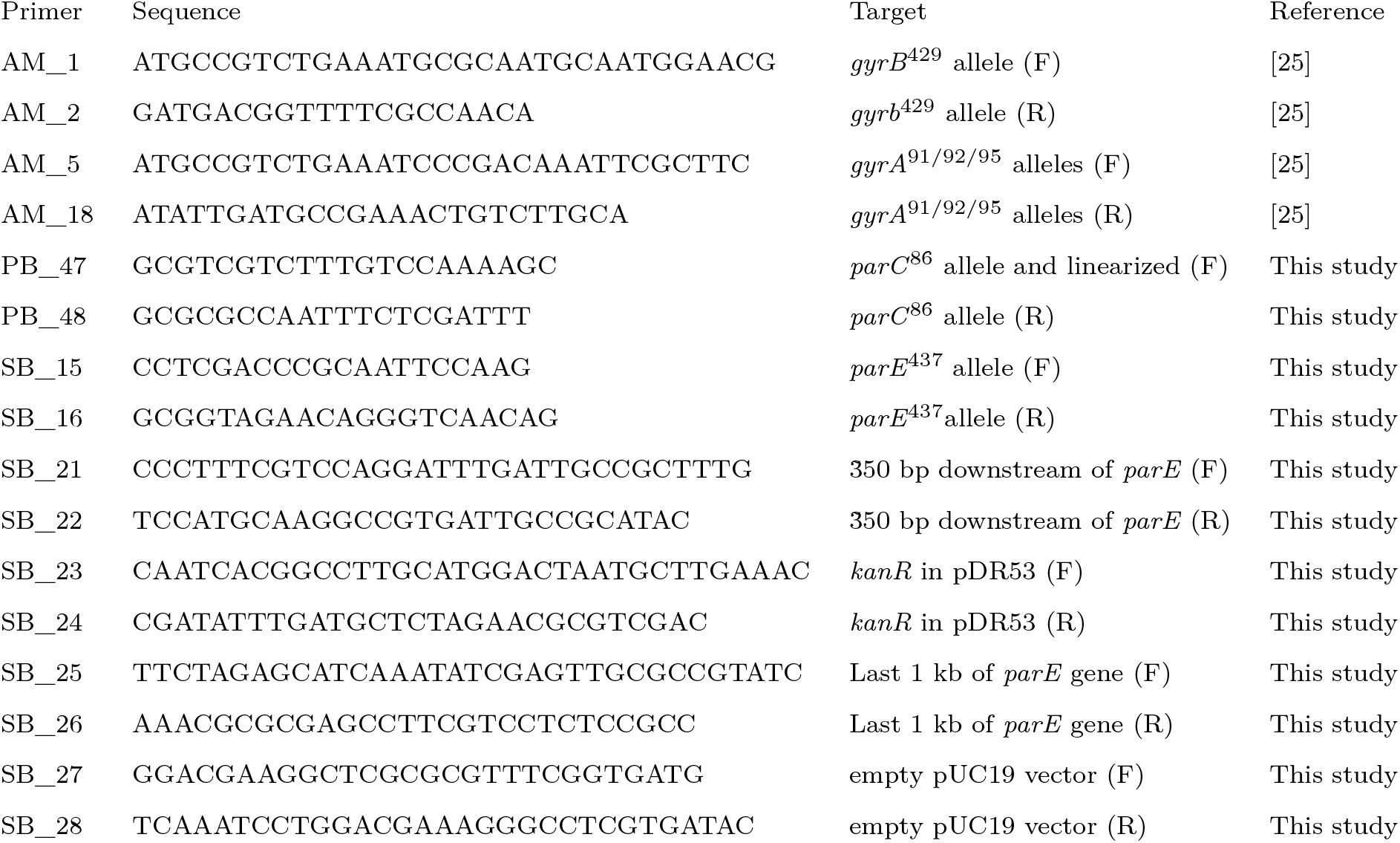
Primers and sequences used in this study.

